# *Insilico* investigation of novel *Plasmodium Falciparum* Glycogen Synthase Kinase(*pf*GSk3β) inhibitors for the treatment of malaria infection

**DOI:** 10.1101/2025.06.07.658456

**Authors:** Kassim F. Adebambo, Sara Ottify

## Abstract

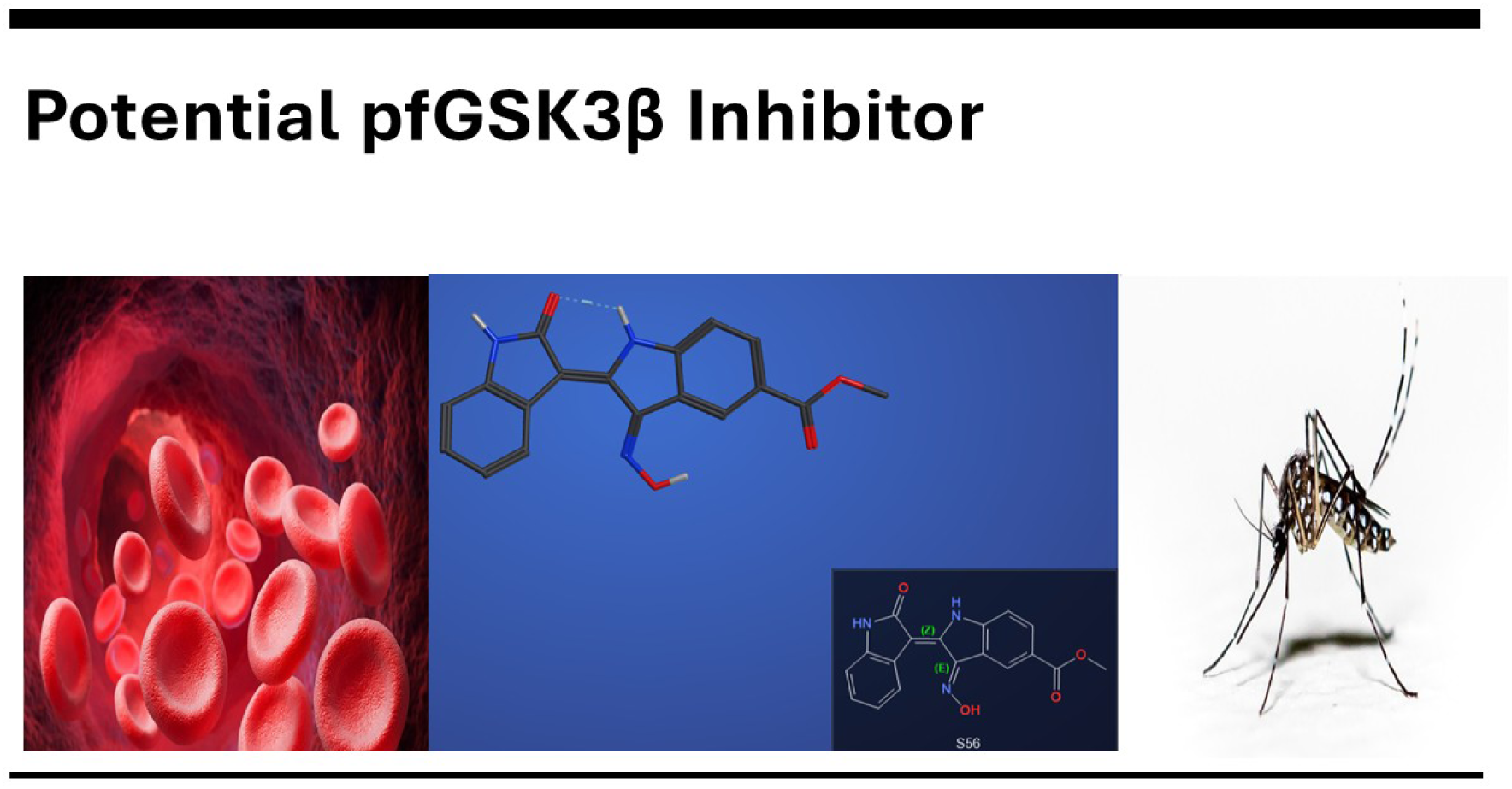

Malaria, a parasitic disease, remains a global health burden with over 260 million cases worldwide in 2023. With the increasing resistance to existing antimalarials, there is an increasing demand in the continuous search for novel therapeutic targets and strategies. Glycogen synthase kinase (pfGSKβ) is a key enzyme involved in metabolic processes such as glycogen synthase phosphorylation, cell differentiation and cell proliferation of the malaria parasite. An attempt to exploit this enzyme as a potential target for drug repurposing was investigated. The insilico method investigated in this research work involved the use of Python coding to mine data from the CHEMBL database for possible GSK3β inhibitors. The data mining using Python coding returned 53 potential molecules ranked according to their 1C50 value. The in silico analysis of these compounds using molecular docking, molecular dynamics simulations showed that three new molecules, labelled **S20** –CHEMBL ID 1910196 (4-[5-(6- hydroxy-1H-indol-2-yl)pyridin-3-yl]benzonitrile), **S39**- CHEMBL ID 2321945- (2-(7-bromo- 2-hydroxy-1H-indol-3-yl)-3-oxoindole-6-carboxylic acid) and **S56**-CHEMBL ID 2321951 (methyl 2-(2-hydroxy-1H-indol-3-yl)-3-nitroso-1H-indole-5-carboxylate), are potential molecules that can be used to inhibit pfGSK3β, of all these three molecules compound **S56** were found to behave better insilico than **S1**- (3,6-diamino-4-(2-chlorophenyl)thieno[2,3- b]pyridine-2,5-dicarbonitrile)-the co-crystallised ligand in pfGSK3β which was used as a control for the insilico drug repurposing research because it has already been tested to inhibit pfGSK3β, Finally, Protox III-an online webserver tool for insilico toxicity study was used to give the potential toxicity of the molecules, this will serve as a tool to guide further research in toxicity of the repurposed molecules. In conclusion, we have been able to propose that three new drug molecules, **S39**, **S20** and **S56,** have the potential of being repurposed for the treatment of Malaria infection.

## Introduction

Malaria is a disease caused by a parasitic protozoan called *Plasmodium*. There are four common types of *Plasmodium*: *P. Falciparum*, *P. Vivax*, *P. Ovale* and *P. Malaria*. The most common type is *Plasmodium Falciparum*. It invades a high number of red blood cells, making malaria a severe and life-threatening disease (Lalloo et al., 2016). Hassan et al (2022 suggested that malaria could be considered as an inflammatory disease because the Malaria parasite can elicit an inflammatory response similar to the inflammatory responses observed in other pathogenic diseases. The malaria parasite is transmitted to humans by infected female mosquitoes (*Anopheles)*. It is distributed in approximately 83 countries and predominantly in the tropical and subtropical areas, including Africa, Asia, Central & south America, parts of the Middle East and some specific islands. Worldwide in 2023, there were over 260 million cases of malaria and over 590,000 deaths, with 76% of those being children under 5 years of age (Travel Health Pro, 2018). Temperatures in the range of 20-30 degrees Celsius, high humidity and seasonal rainfall, forest ecosystems, as well as rural areas, all increase the risk of exposure to malaria. Even though there is a higher risk of being exposed to mosquito bites at Dawn and Dusk, the risk of being bitten is always present at any time during the day, whether indoors or outdoors (UK Health Security Agency, 2014). Lacerda-Queiroz et al, 2011 reported that a single infection of malaria can lead to high morbidity and fatality if not controlled.

### Treatment/Resistance

There are important factors to consider when initiating treatment. For instance, it is important to determine the species of *Plasmodium* for the treatment of malaria, this is because species such as *P. falciparum* may rapidly progress to cause a life-threatening condition, while other species may be less severe. Furthermore, *P. vivax* and *P. Ovale* will require treatment to target the dormant hypnozoites that relapse later. Depending on the symptom presented, malaria will be classified as uncomplicated or severe, which will then determine treatment. Establishing the likelihood of drug resistance and its patterns stems from the knowledge of the geographical area that is acquired. This helps to determine the appropriate treatment or combination of treatments. However, Malaria parasite has been found to rapidly develop resistance to the common drugs that are currently being administered in the clinic, for example, Malaria has found a way to rapidly develop resistance to chloroquine and other drug like artemisinin (Li et al, 2016), shockingly, Nicholas J white in his report in 2004 stated that while Malaria infection has developed resistance to most of the frontline drug our hope was in Artemisinin and he concluded that if should lose Artemisinin to resistance we are going to face a problem of having untreatable malaria, but Li et al 2016 has shown that Malaria now show resistance to Artemisinin, therefore there is now an urgent need to investigate a novel target and novel drug molecules for the treatment of Malaria infection.

### Glycogen Synthase Kinase and its role in malaria

Glycogen synthase kinase-3 (GSK-3β) is a serine/threonine protein kinase that consists of two isoforms. it was found to phosphorylate glycogen synthase but is also involved in many biological processes (Eldar-Finkelman, 2002), such as cell proliferation, differentiation and protein synthesis (Masch & Kunick, 2015). Serine is a non-essential amino acid involved in cell growth and in the synthesis of purine, adenine and guanine bases in DNA (Drugbank, 2005a). Threonine is an amino acid that helps in fat metabolism in the liver (Drugbank, 2005b). It was found that GSK3β, along with other protein kinases, are crucial for the parasite’s ability to proliferate in red blood cells. However, in *P. falciparum*, pfGSK-3β is critical for schizogony, and in infected red blood cells, it undergoes phosphorylation at tyrosine 229. Due to the hypothesis that GSK-3β in *P. falciparum* engages in roles such as cell cycle control, differentiation process and metabolism regulation, it is seen as a potential drug target to overcome resistance against antimalarial drugs (Masch & Kunick, 2015). The malaria P. falciparum genomic data have been found to contain about 65 eukaryotic protein kinases (Ward et al, 2004); some of these kinases are useful in the process of host cell infection of the malaria parasite(Hitz et al, 2021). As of today, only one antimalarial drug has been successfully developed to target the kinase enzyme. This drug targets the PI4K enzyme; this drug is highly tolerable and has a very high anti-malaria property. currently, the PI4K inhibitor is still at the clinical trial stage (Sinxadi et al, 2020 and McCarthy et al, 2020). Furthermore, the other enzyme that has been studied is pfGSK3β, this research work is based on investigating novel inhibitors for pfGSK3β.

### Target selection

GSK-3β in *Plasmodium Falciparum* (pfGSK3β) has been observed to help in the parasite’s survival by altering red blood cell metabolism, membrane transport and cytoskeletal properties, possibly increasing its growth. The protein kinase may also help to upregulate conductive and new permeation pathways. Subsequently, it has been detected that interference with circadian clock regulators could assist in the maturation and replication of blood stages of parasites (Masch & Kunick, 2015); therefore, targeting this protein could be an effective treatment against malaria infection. The 3D structure of pfGSK-3β is shown below in Figure 1, and the various molecules synthesised and tested by Masch and Kunick et al, 2015 as pfGSK3β inhibitors are shown in Figure 2.

**Figure 1.**
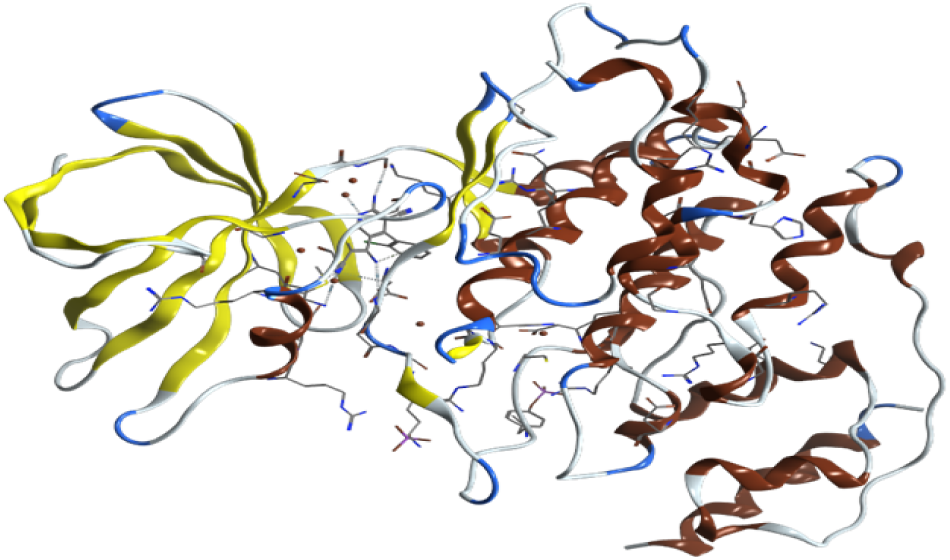
3D structure of Plasmodium Falciparum Glycogen Synthase-3 PDB ID: 3ZDI

**Figure 2:**
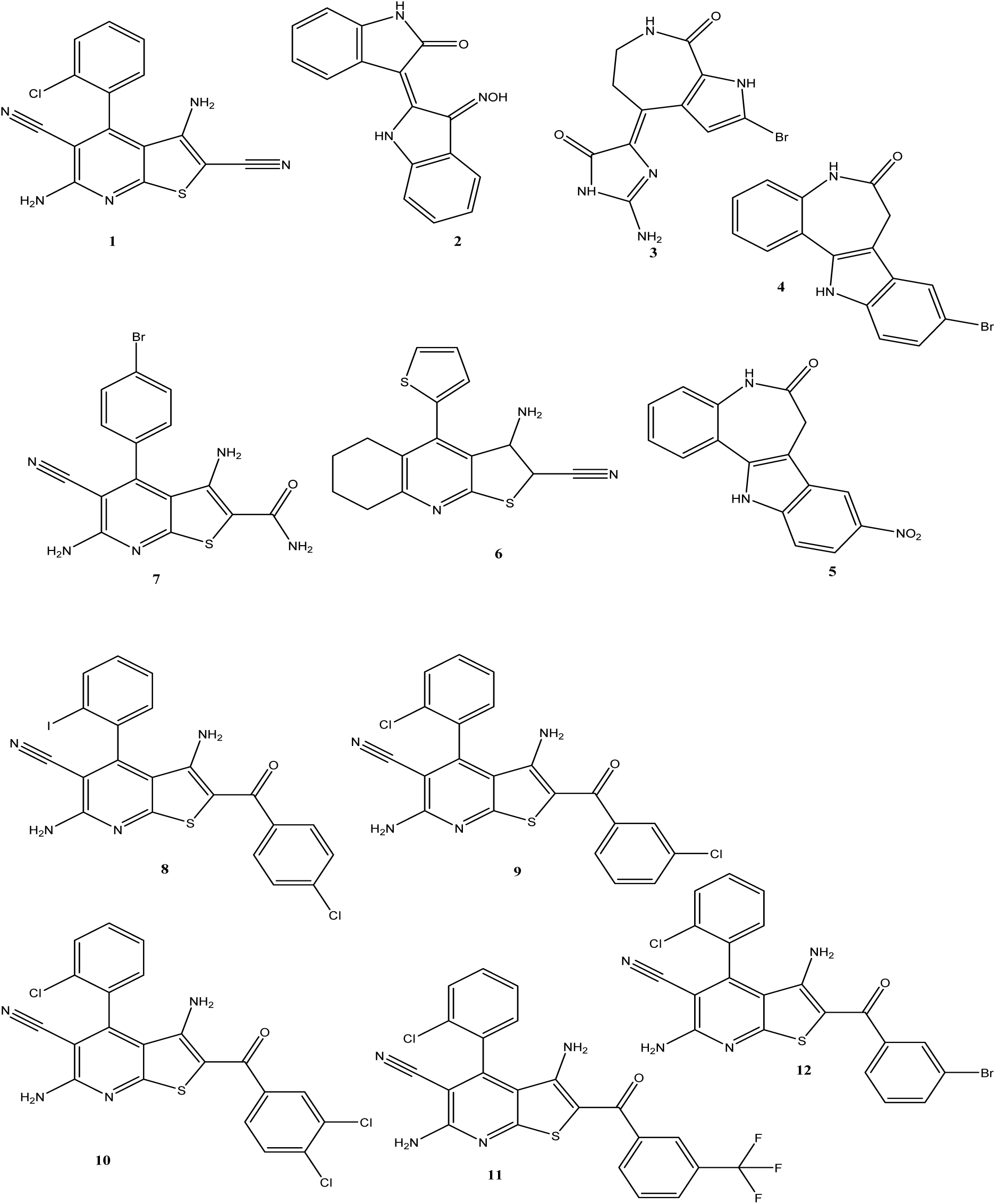
Possible drug molecules that have been investigated for the inhibition of pfGSK3β by Masch *et al*, 2015

### Challenges in malaria management

As mentioned above, malaria is one of the most life-threatening diseases, making it a significant health burden. There are multiple barriers to the successful management of the disease. This includes the resistance to anti-malarial drugs such as chloroquine, the use of similar derivatives of chemical compounds contributing to cross-resistance and genetic mutations within parasites.(Kokwaro, 2009). These challenges highlight the necessity for ongoing research into the development of novel drugs and therapeutic targets for the effective treatment of malaria infection.

### Aim of the Research

This research was inspired by the work of Masch et al 2015, where they started working on developing a new drug molecule for the inhibition of a new drug target in the Malaria parasite, the pfGSK3β. Most of the compounds synthesised and tested by Masch et al, 2015 are shown in Figure 2, and these molecules serve as a template for our insilico investigation. The application of computational studies is now becoming more important in the era of machine Learning to speed up the development of novel drug molecules. Therefore, using the potential drug molecules and the targets reported by Masch et al, 2015 as a control, we investigated the possibility of using computational and data science tools like Python to repurpose the already synthesised and tested GSK3β inhibitors reported in the CHEMBL database as pfGSK3β inhibitors. This insilico approach is very important because of the high risk, cost, environmental impact and time factor that goes into the development of a new drug from scratch. Our approach has been justified by Wu et al, 2019 and Parvantaneni, 2019. These two groups have reported the various successes that have been recorded by the insilico research. Furthermore, Computational investigation has played an important role in the repositioning of the existing biomolecules to the treatment of new diseases, for example the use of computational investigations has been exploited which involved a scoring and ranking model (Arul et al, 2022), Structure-based drug repurposing research using insilico tools has been reported by Choudhury et al in 2022.

Finally, we aimed to use Python to generate code for the search of existing GSK-3β inhibitors from ChEMBL database, investigate the binding affinity of the bioactive molecules obtained from the Python search in pfGSK3β, using PDB 3ZDI, to validate this drug repositioning study, the binding affinity will be compared with that of the molecules in Figure 2. Furthermore, Molecular Dynamics Simulations of the potential drug compounds will be investigated using Gromacs tools as well as the insilico toxicity prediction, using a web server insilico toxicity study known as Protox III. The data obtained from these computational studies will be used to conclude whether a novel potential drug molecule has been discovered for the inhibition of pfGSK3β or not.

## Methodology

### Data Mining Study on CHEMBL Database Using Python Coding

Python is a programming language for data science and machine learning (Gayathri Rajagopalan, 2021). The Python tool has become a very useful tool in drug repurposing because it enables scientists to mine a big data bank for drug repositioning studies.. The Python used in this research was launched from the Anaconda platform. Anaconda-Navigator was installed using this weblink: https://www.anaconda.com/download/success, The ChEMBL database is an open database containing many drugs, like bio-active compounds, with their biological activities, functional and ADMET information (Gaulton et al., 2011).

According to Michal Nwotka 2015, Python software has been developed in order search the CHEMBL database for data mining, This Python tool is easy to use because the CHEMBL database has created a client library API for easy data mining on the CHEMBL database.

A typical data mining procedure that was modified by our team is given below

*from chembl_webresource_client.new_client \import new_client*

*# Receptor protein-tyrosine kinase erbB-2chembl_id = "CHEMBL1824" activities = new_client.mechanism\. filter(target_chembl_id=chembl_id)*

*compound_ids = [x[’molecule_chembl_id’] for x in activities]*

*approved_drugs = new_client.molecule\. filter(molecule_chembl_id in=compound_ids)\. filter(max_phase=4)*

https://www.researchgate.net/publication/304802966_Want_Drugs_Use_Python *[accessed May 29 2025]*.

Starting from the above coding step, we compiled codes which enable us to mine for all GSK3β inhibitors in the CHEMBL database. The coding flow chart we developed is represented in Figure 3 below. The data mining returned 53 possible inhibitors.

**Figure 3:**
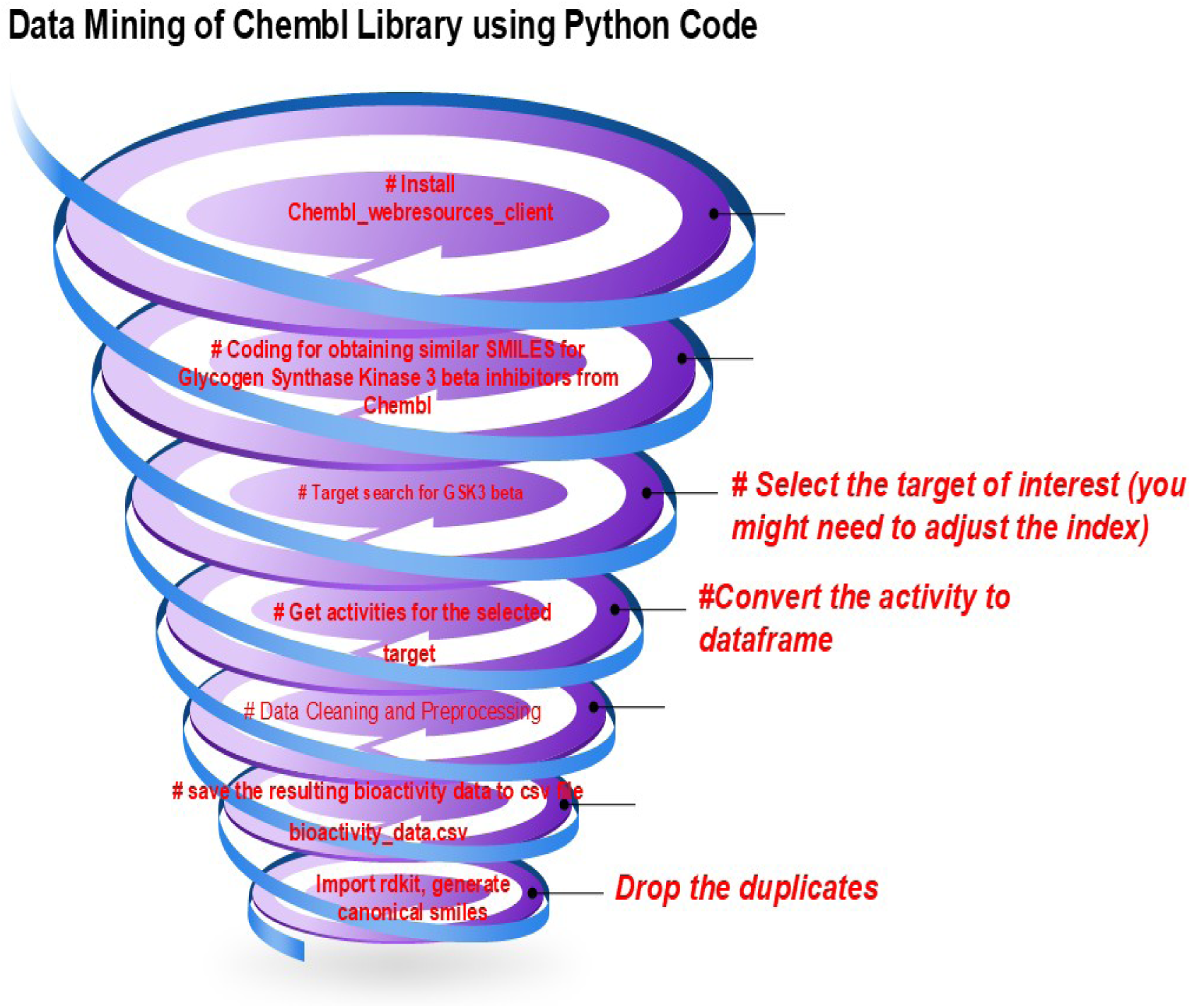
Data mining flowchart for obtaining GSK3β inhibitors from the CHEMBL Database

### Molecular docking

Our research was inspired by the work already reported by Masch et al, 2015. In their report, they showed that some molecules were found to inhibit pfGSK3β, a novel target for treating Malaria. We used the designed molecules [Figure 2] as a control for our insilico research. Therefore, 65 compounds were used in the molecular docking experiment, twelve from the Masch et al, 2015 compound and 53 from the Data mining of the CHEMBL library. The molecular docking experiment was carried out using Molecular Operating Environment (MOE) software, a comprehensive software system used by medicinal chemists, biologists, crystallographers and computational chemists (Chemical Computing Group, 2024). The general flowchart for molecular docking, as reported by Morris et al 2008 is shown in Figure 4 below

**Figure 4.**
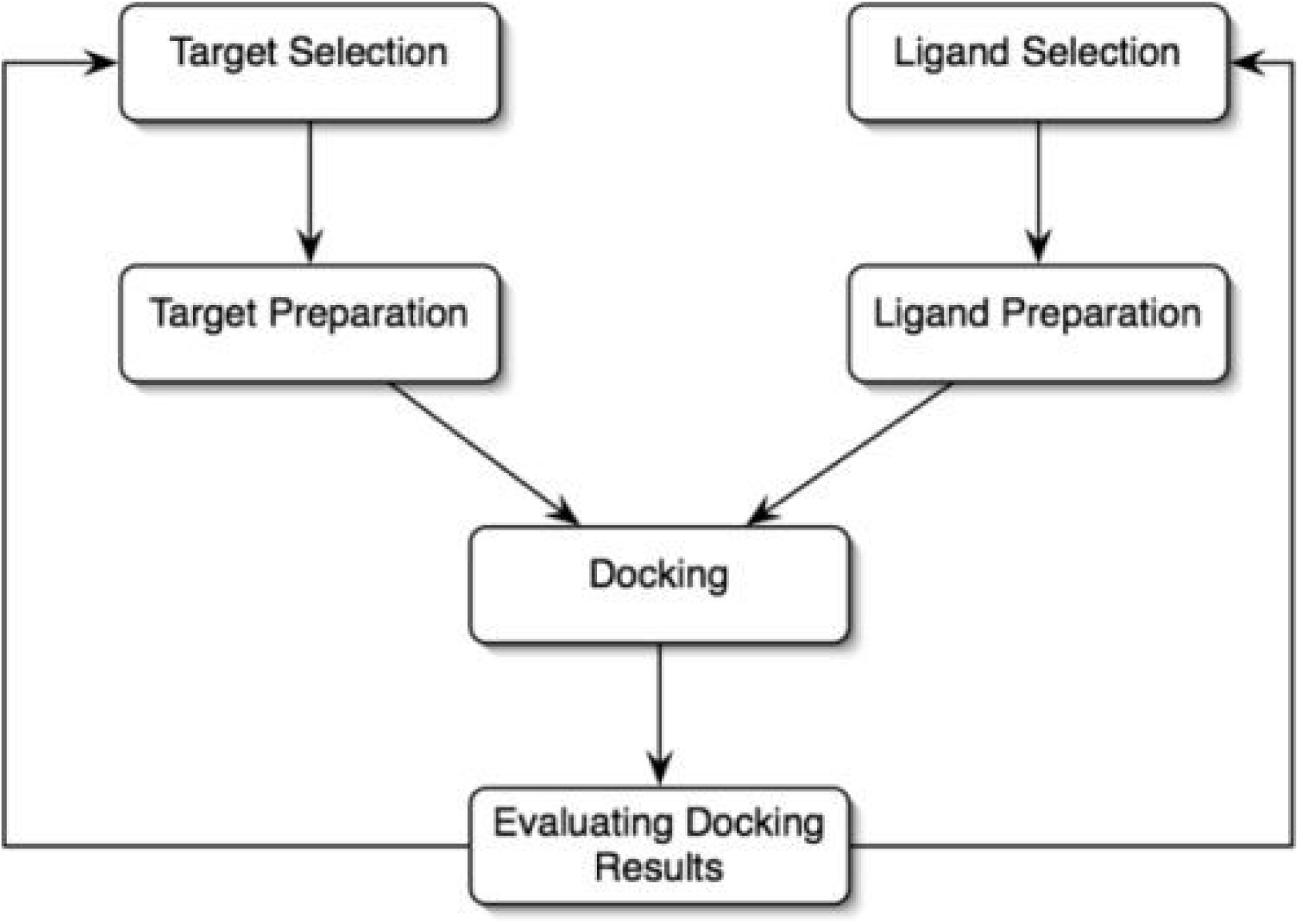
Steps involved in docking. (Morris & Lim-Wilby, 2008).

### Ligand Preparation

As shown in Figure 4, Ligands need to be prepared before carrying out molecular docking study. The ligand preparations is a crucial step in protein-ligand interaction as it involves building the ligands and minimising their energies, thereby making the ligands suitable for efficient protein-ligand interactions. During the preparation, SMILES of the molecules were input into MOE by using the builder command, the final structures were then minimised. The prepared ligand structures are converted into an .mdb file ( a format suitable for MOE docking software)

### Protein Preparation

A 3D copy of the 3ZDI was downloaded from the Protein Data Bank website. The protein was prepared for docking using the quick prep ion of the MOE software. the preparation involved preserving the sequence, neutralising the receptor, and all the water molecules farther than 4.5Å from the ligand or receptor were deleted. Furthermore, all the missing amino acids were fixed. Protein was refined to an RMS gradient of 0.1 kcal/mol/A^2. The site finder icon on MOE was used to determine the binding pocket of the co-crystallised ligand, the pocket served as the target site for the binding of the repurposed drug molecules.

### Molecular Docking- Protein-Ligand Interaction

All 65 prepared molecules were docked into the identified binding pocket of the co-crystallised ligand. For this experiment, a triangular placement matcher was applied, and the docking was done with rigid receptor refinement. Furthermore, 30 binding poses were carried out (London dG-30 poses), and the system was programmed to return the five best poses ( GBVI/WSA dG- 5 poses). Rigid receptor placement means that the receptor/ protein remains fixed while the ligand is flexible to undergo conformational changes, but the protein’s structure does not adjust.

### Molecular Dynamics Simulations study

The Molecular Dynamics simulations were carried out using Gromacs 2023.1 on a GPU, the procedure described by Lemkul et al 2018 on <u>Protein-Ligand Complex</u> was followed step-by-step. The protein 3ZDI obtained from the PDB has an inbuilt amino acid code that needed to be resolved, because this was not part of the forcefield residue data. The error was resolved by using the Protein Repair and Analysis Server (<u>Protein Repair & Analysis Server</u>). The output PDB obtained from the Protein Repair and Analysis Server was used again for the re-docking of the 65 ligands; this was necessary in order to investigate whether the repaired PDB structure would affect the docking scores and ranking of the 65 ligands under investigation. The rescoring did not show any changes in types and number of ligand interactions formed by the 65 molecules. Therefore, this gave confidence that the repair did not have any impact on our docking experiment. The molecular dynamics simulations run involved the use of the CHARMM all-atom force field to build the protein topology, and the official general force field server CGenFF to build the ligand topology. Molecular Dynamics Simulations production was run for 100ns.

### Insilico Toxicity Study

Drug development involves testing potential analogues for their safety. One of the criteria for drug safety is its toxicity level. taking a potential drug molecule from the laboratory to the market involves a huge amount of investments, and many drug compounds never make it past the toxicity test (Guengerich et al, 2011). Most drugs failed pre-clinical tests invivo and in vitro, therefore, it is a good study if in silico-preclinical trials could be used to screen most drugs for their potential toxicity before investing in their invivo study. This saves time and reduces the environmental stress as a result of wet laboratory experiments. The use of Protox III has been developed to support clinical research. Therefore, our Insilico toxicity study was carried out by using a web server service <u>ProTox-3.0 - Prediction of TOXicity of chemicals</u>, (Pryanka Banerje et l, 2024). The online tool was used to carry out insilico analysis of the potential drug candidates investigated in this research.

## Results and Discussion

This research aimed to use Python coding to mine the CHEMBL library for possible GSK3β inhibitors, which we could repurpose towards the inhibition of pfGSK3β. The coding was successfully designed; this data mining generated 53 molecules after ranking them according to their IC50 (Table 1)

**Table 1:**
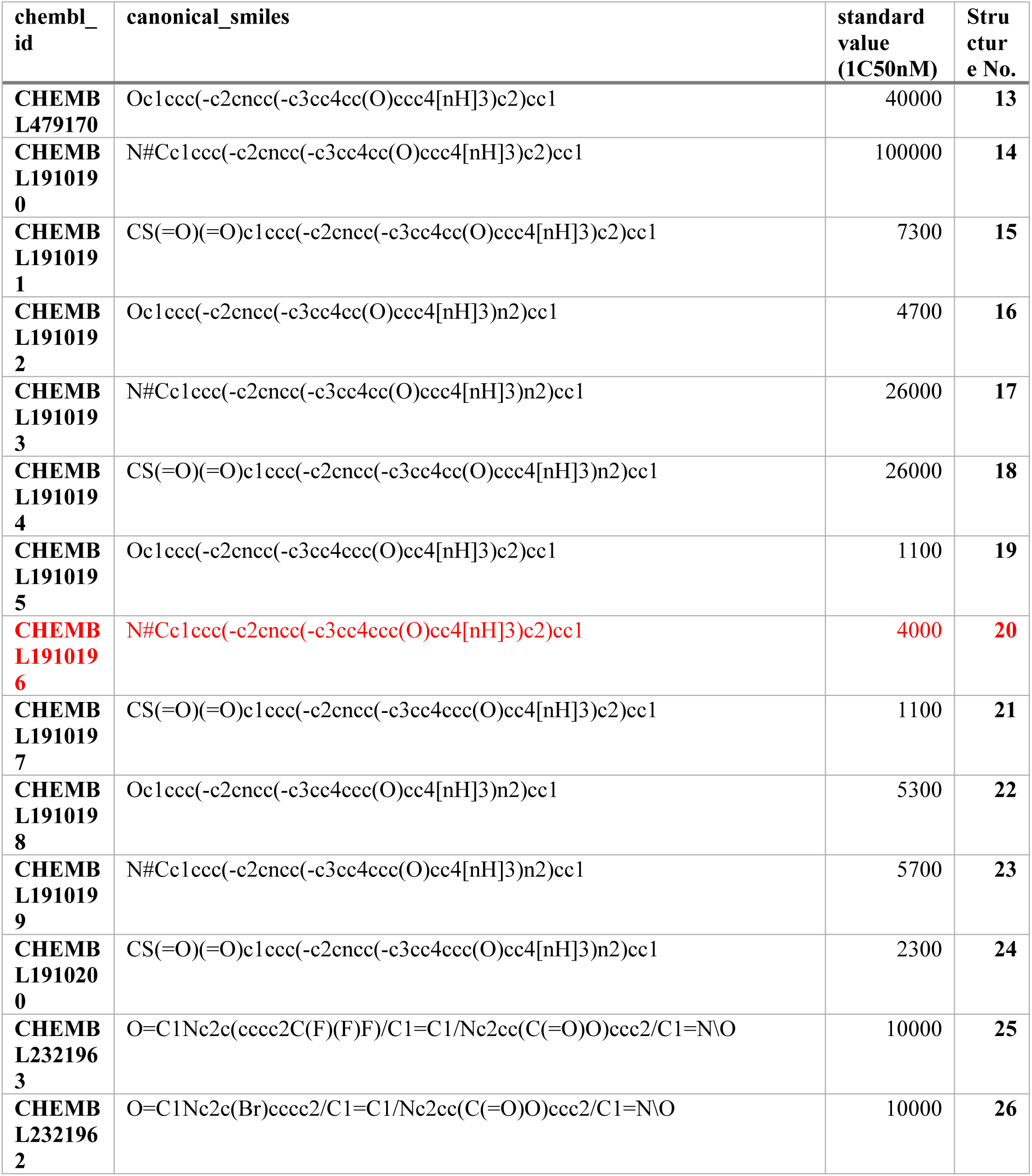
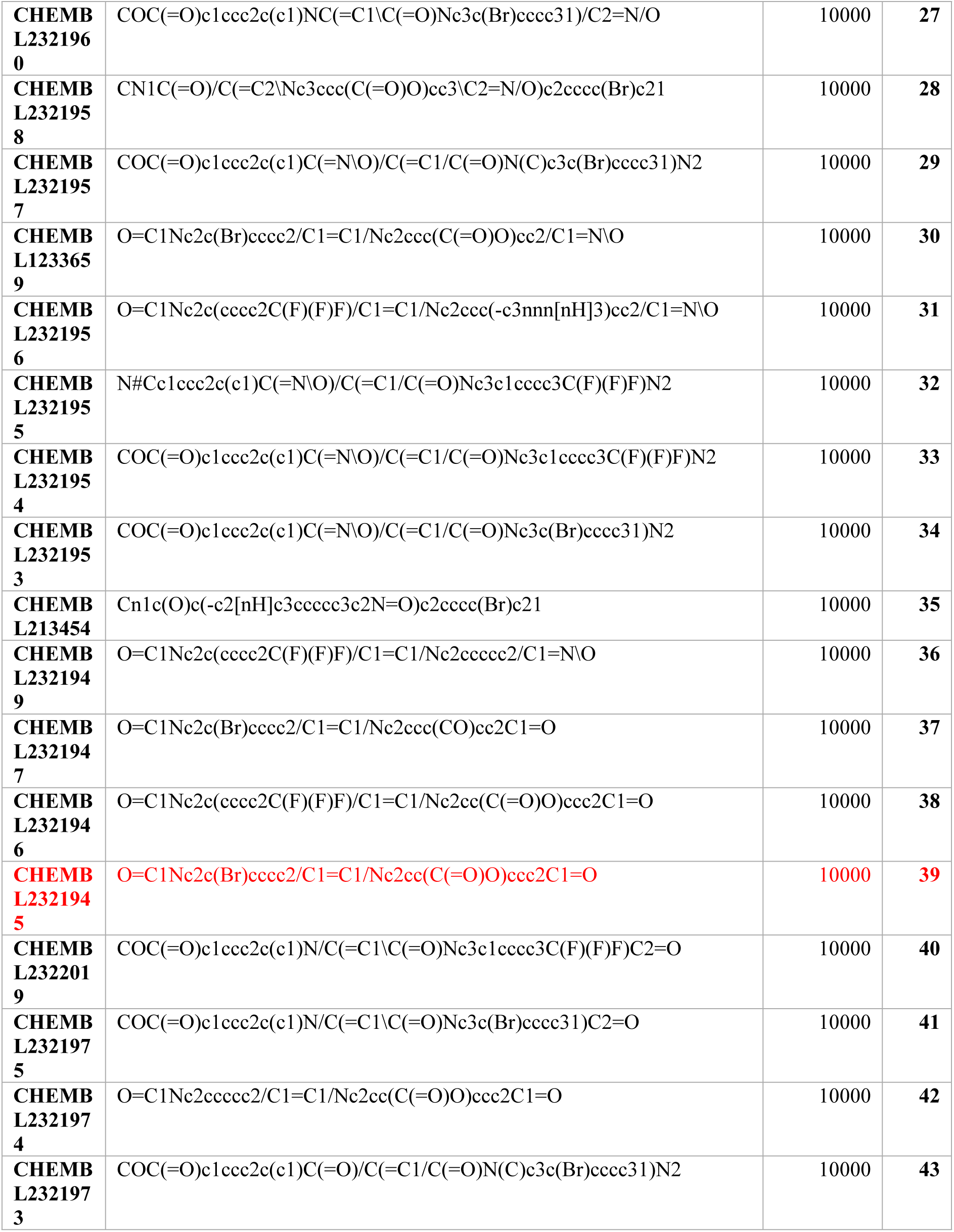
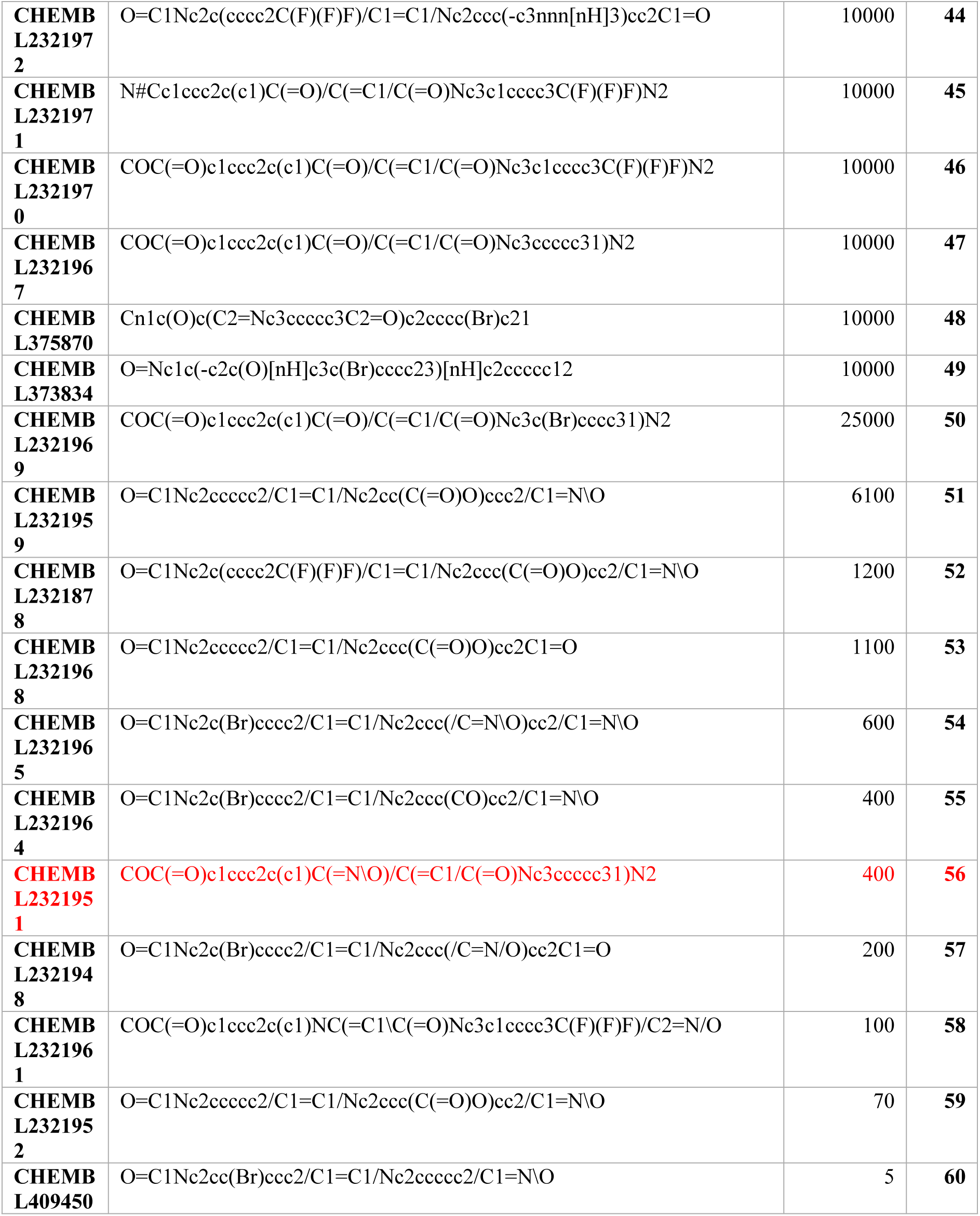
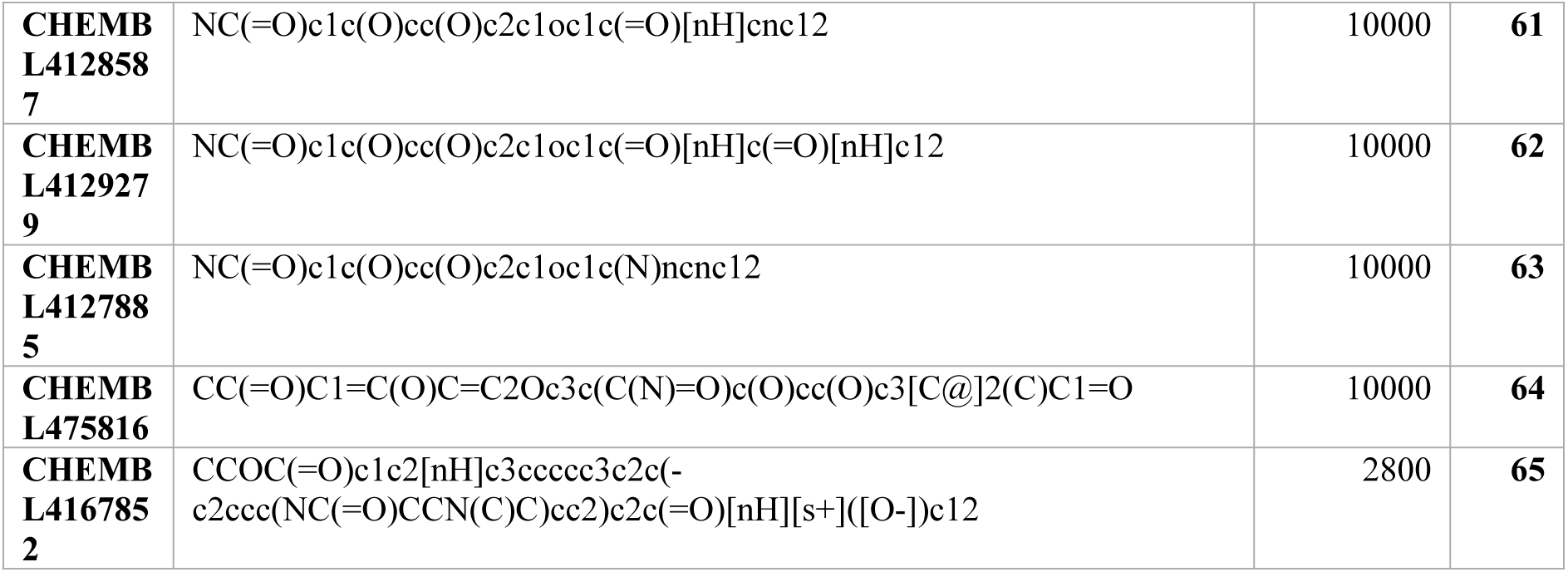
53 molecules generated from the Data mining of the CHEMBL database using the generated coding.

### Molecular Docking Investigation

The initial step in the molecular docking was to first obtain the SMILES of the 12 structures obtained from the Masch et al 2015 paper using ChemDraw software. Furthermore, the 3D structure of pfGSK3β (Figure 5A) was downloaded from the protein databank, which contains a co-crystallised ligand (in this research, the ligand will be referred to as **S1**). The position of the co-crystallised ligand in the receptor enables us to identify the binding site of the receptor. (Figure 5B). To understand the shape of the binding pocket, a surface mapping study was carried out on MOE software, the surface mapping revealed the shape of the binding pocket Figure 5C.

**Figure 5:**
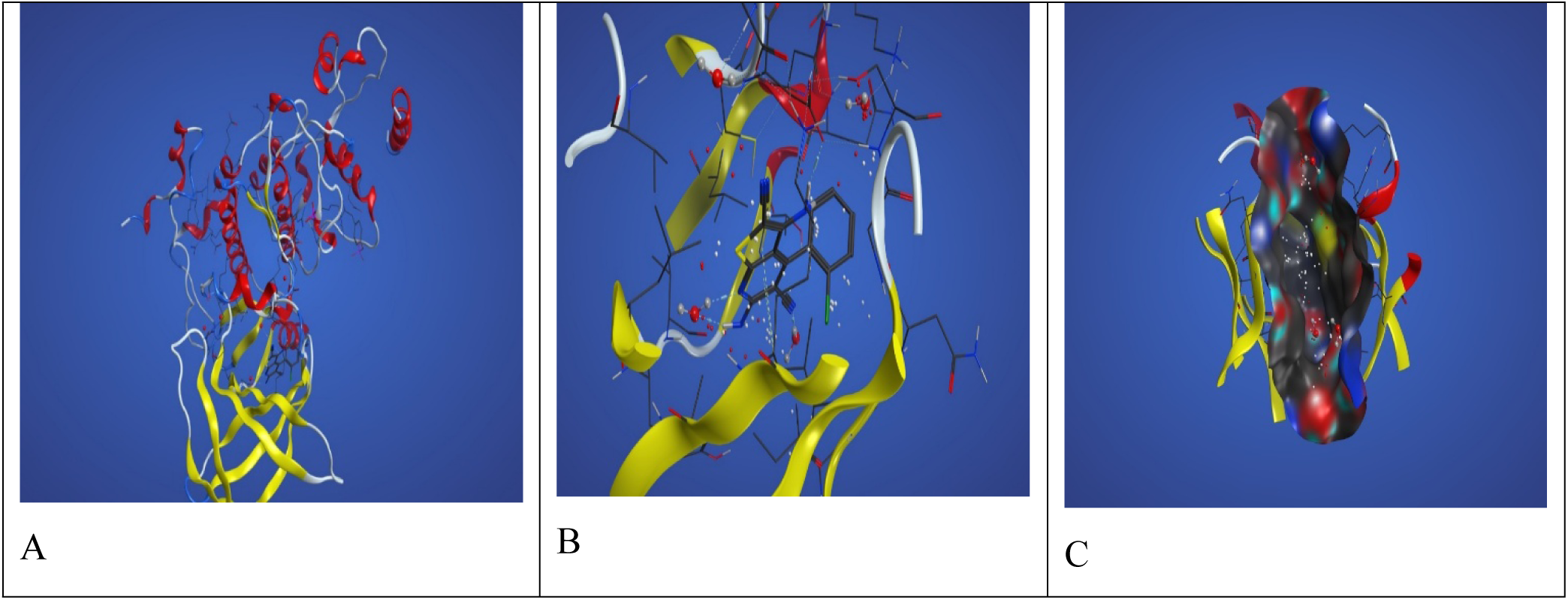
A. Structure of 3ZDI embedding the co-crystallised ligand. B. Co-crystallised ligand in the binding pocket, and C is the surface mapping of the binding pocket.

Molecular docking is a computational tool which allows us to model interactions between ligands and proteins based on the lock and key theory (Figure 6). Through this application, we can predict the binding capacity of molecules (ligands) to their protein (receptor) and explore how the ligand could be coupled in the conformational space (Silva et al., 2019), with the main aim being to find the most stable receptor-ligand complex through their binding affinities.

**Figure 6.**
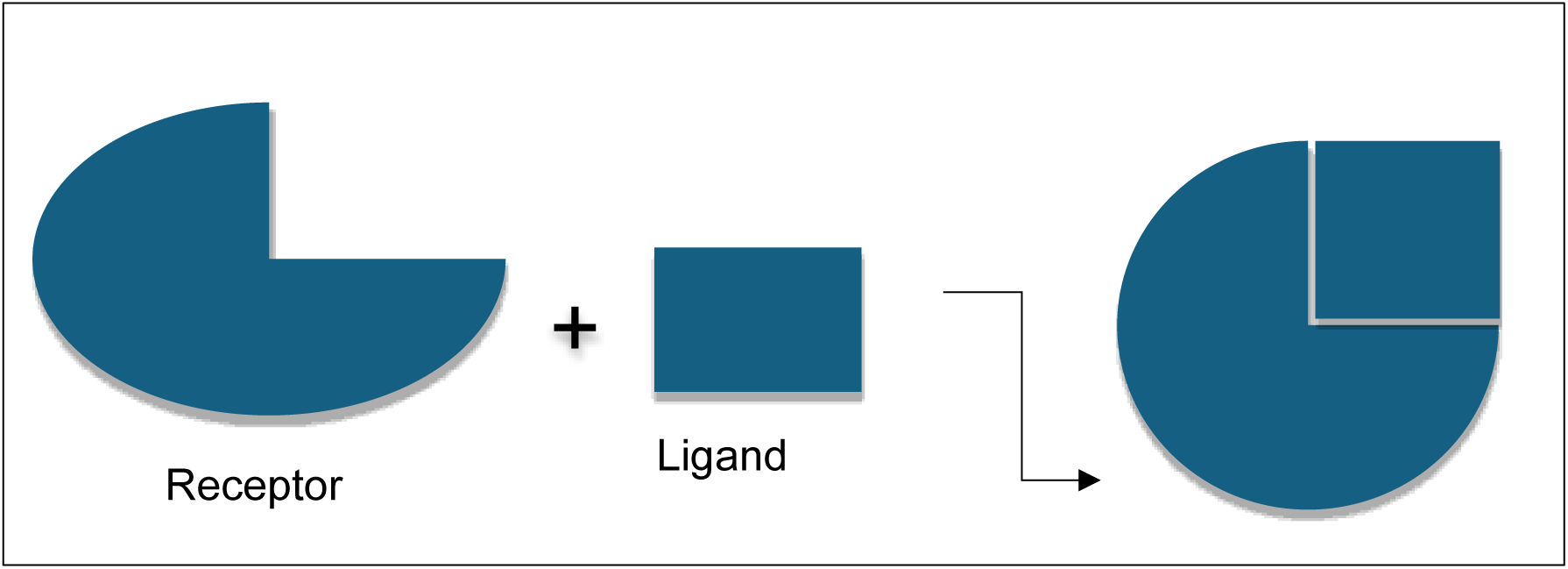
Lock and key theory (Tripathi et al, 2017).

### Ligand-Receptor Interactions

The following ranking system was used to determine which Ligand would be best suited for the receptor.

1. Molecules with a high number of interactions, containing both hydrophilic and hydrophobic interactions, with a very high binding energy will be rated the best. Hydrophobic interactions, such as pi-pi bond, will provide thermodynamic stability as well as increase the tendency of forming contacts among themselves rather than with polar water (Ferenczy & Kellermayer, 2022). Whereas hydrophilic interactions, such as hydrogen bond acceptors/donors, can enhance binding specificity and affinity between the ligand and the protein binding pocket (Dinesh Kumar Sriramulu & Lee, 2020).
2. Low binding energy with a high number of interactions, with the presence of both electrophilic and hydrophobic interactions, was rated second
3. Molecules with no interactions were not considered.

Using the above ranking criteria, the best four ranked molecules from the molecular docking analysis are given in Table 2 below. For ease of discussion, the following key words will be used. The IUPAC names were confirmed using the PubChem search (https://pubchem.ncbi.nlm.nih.gov/)

**Table 2:**
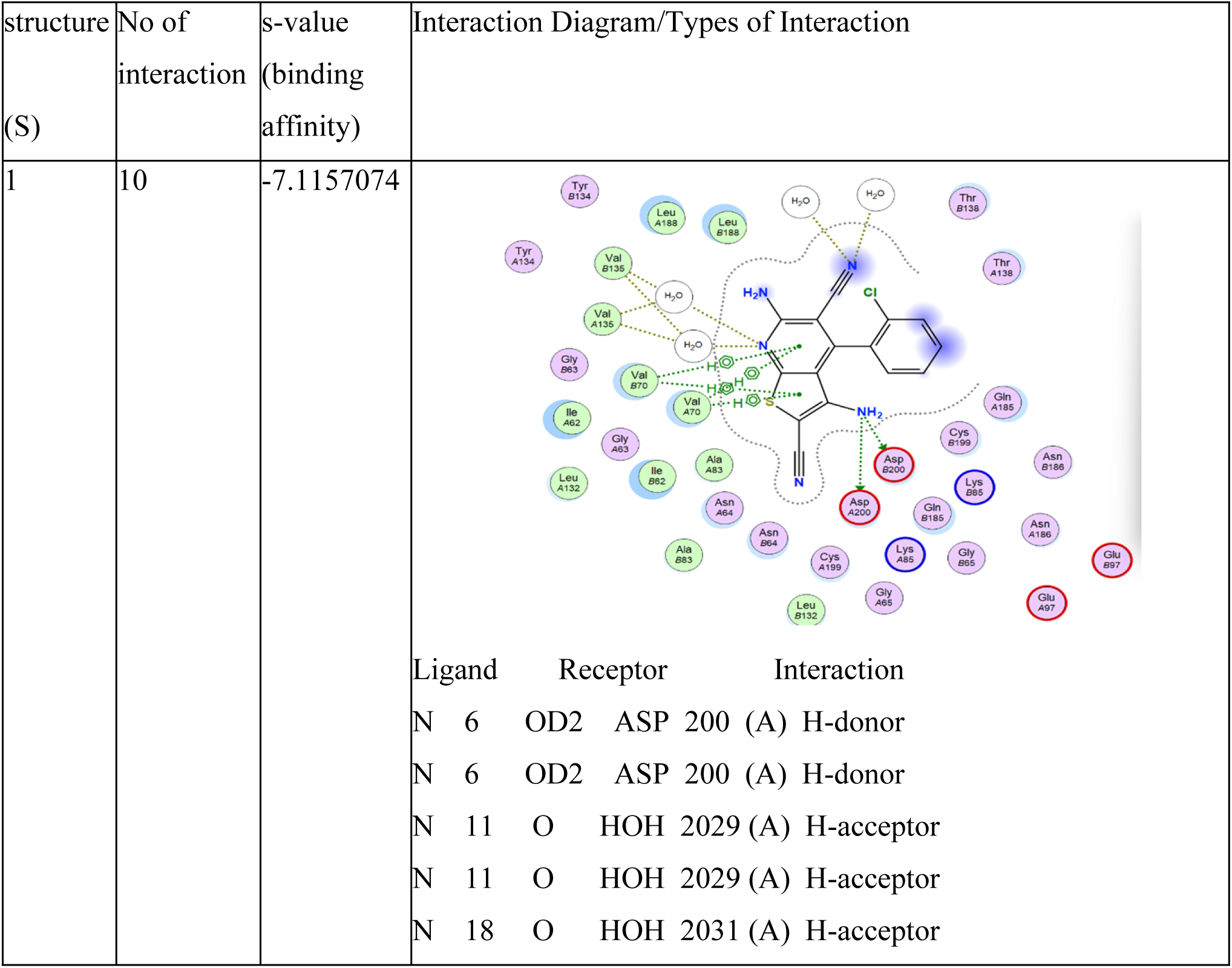
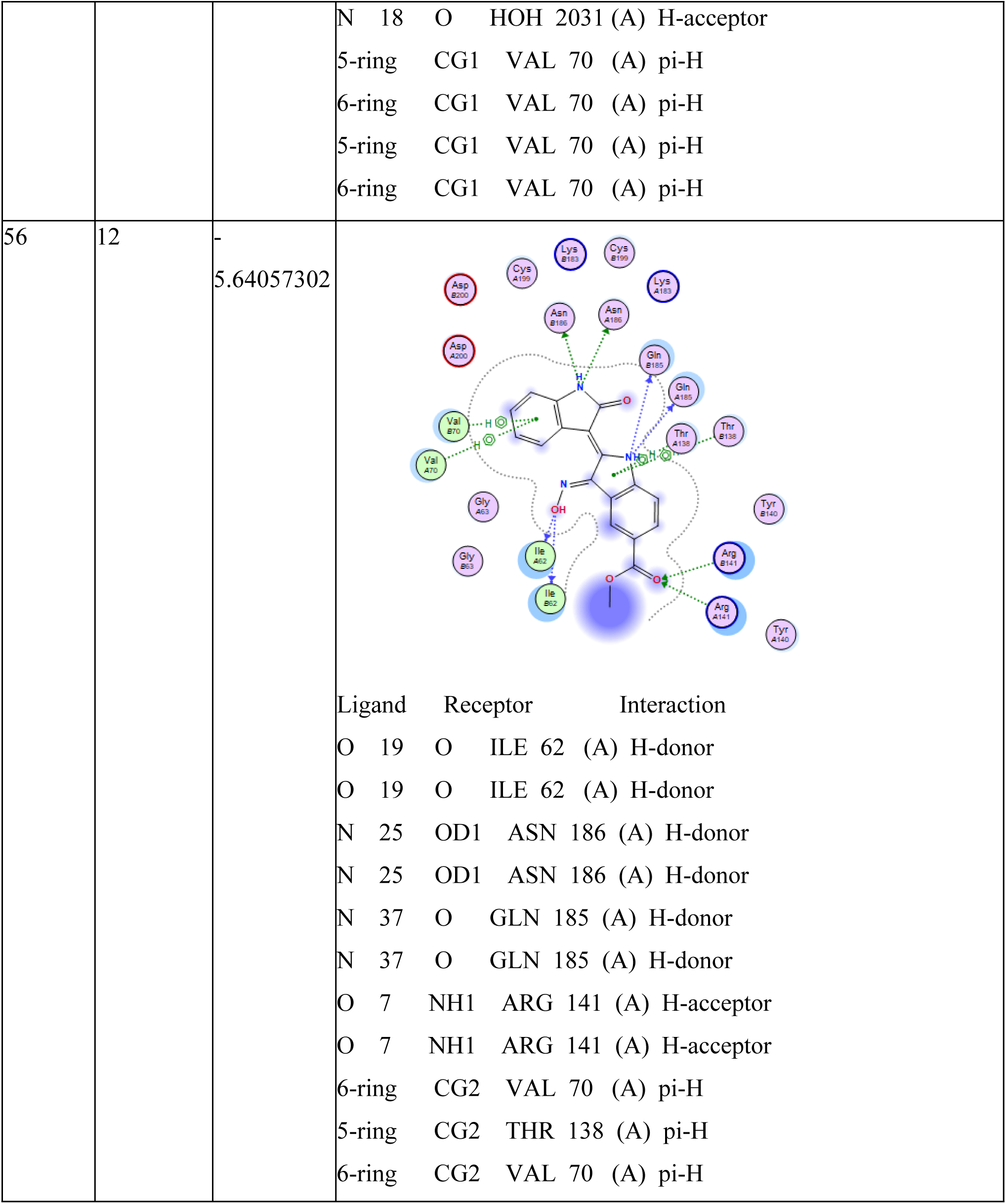
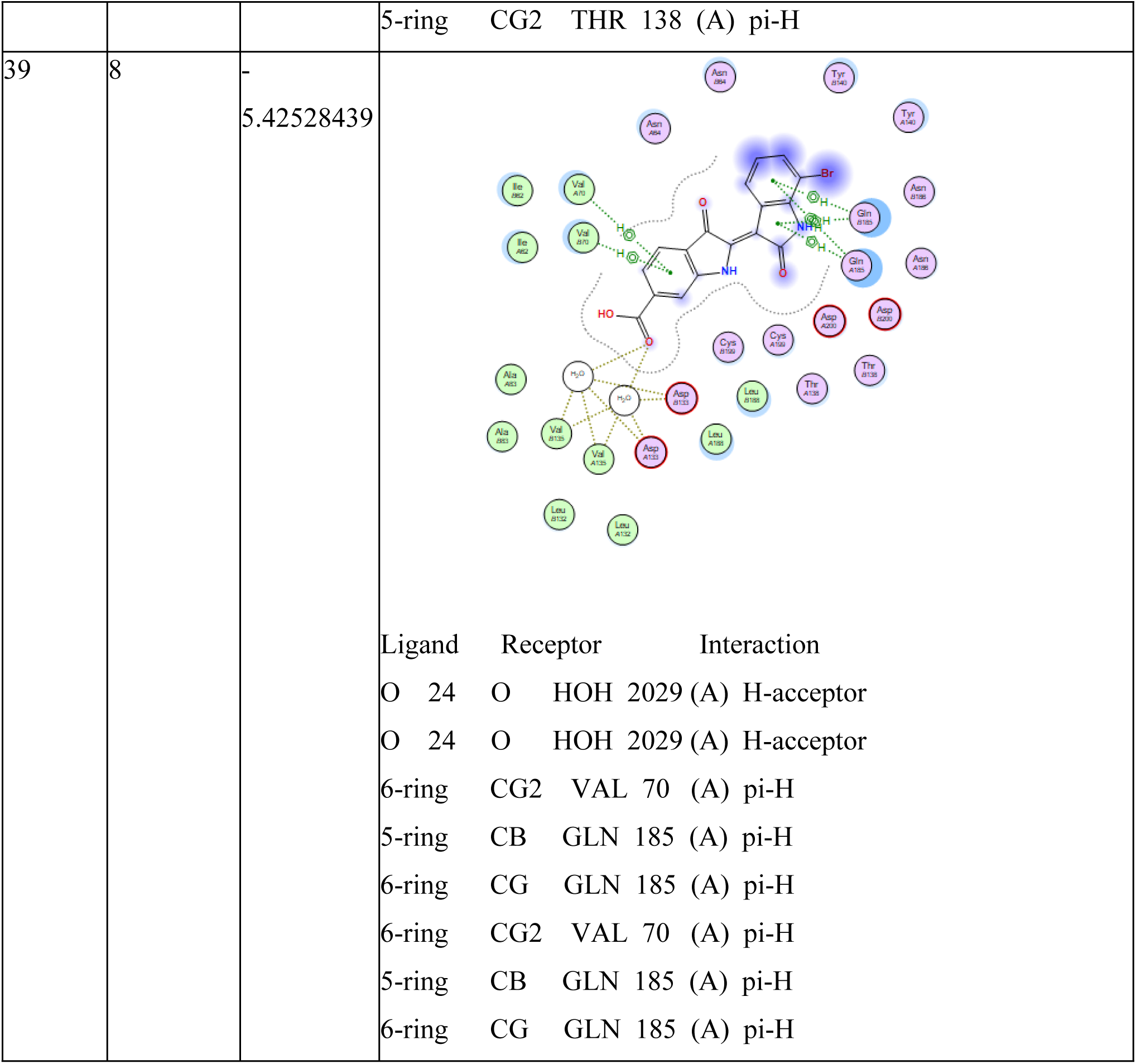
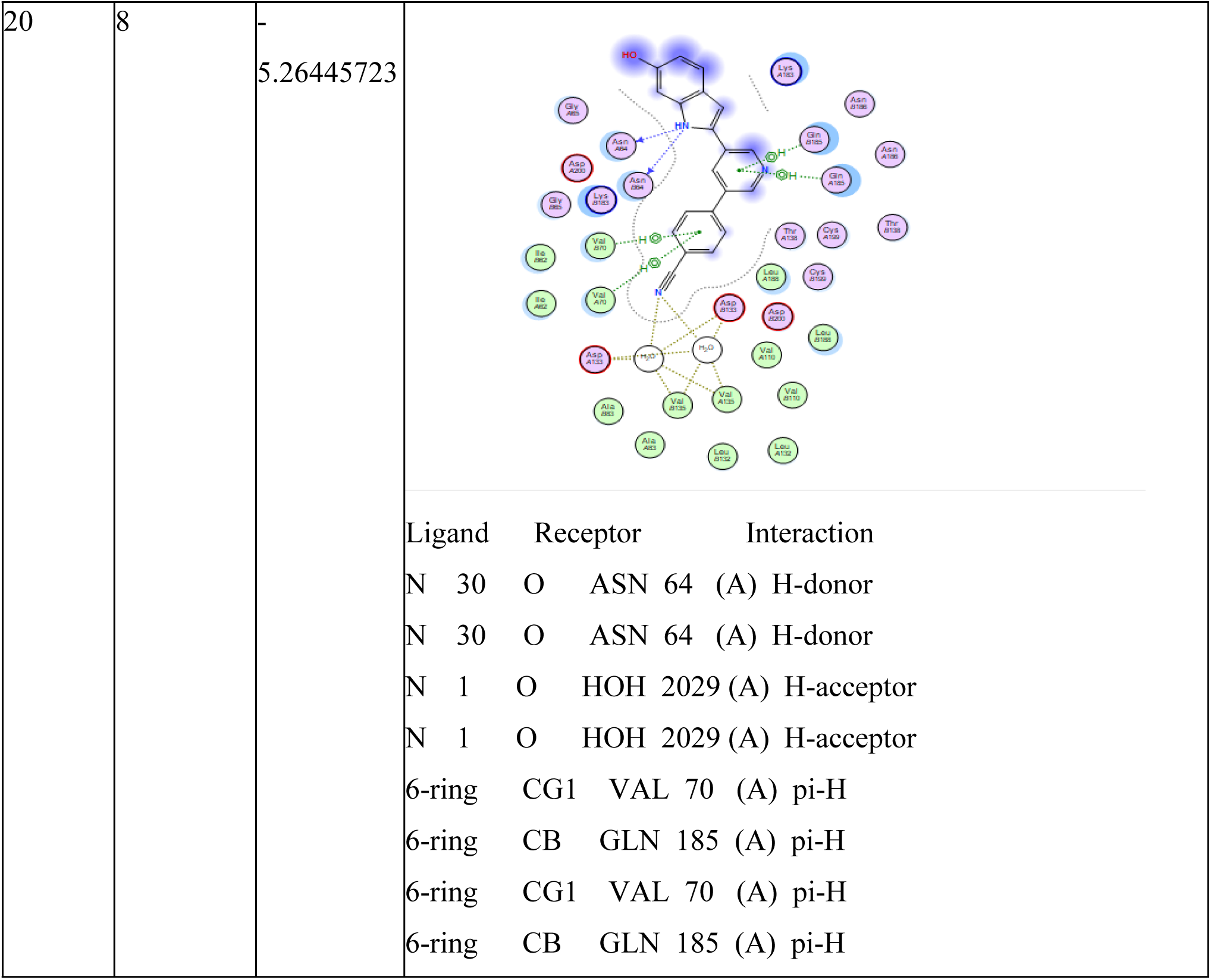
Protein Ligand Interactions of the four best-ranked molecules obtained during the molecular docking investigation.

**Table 3:**
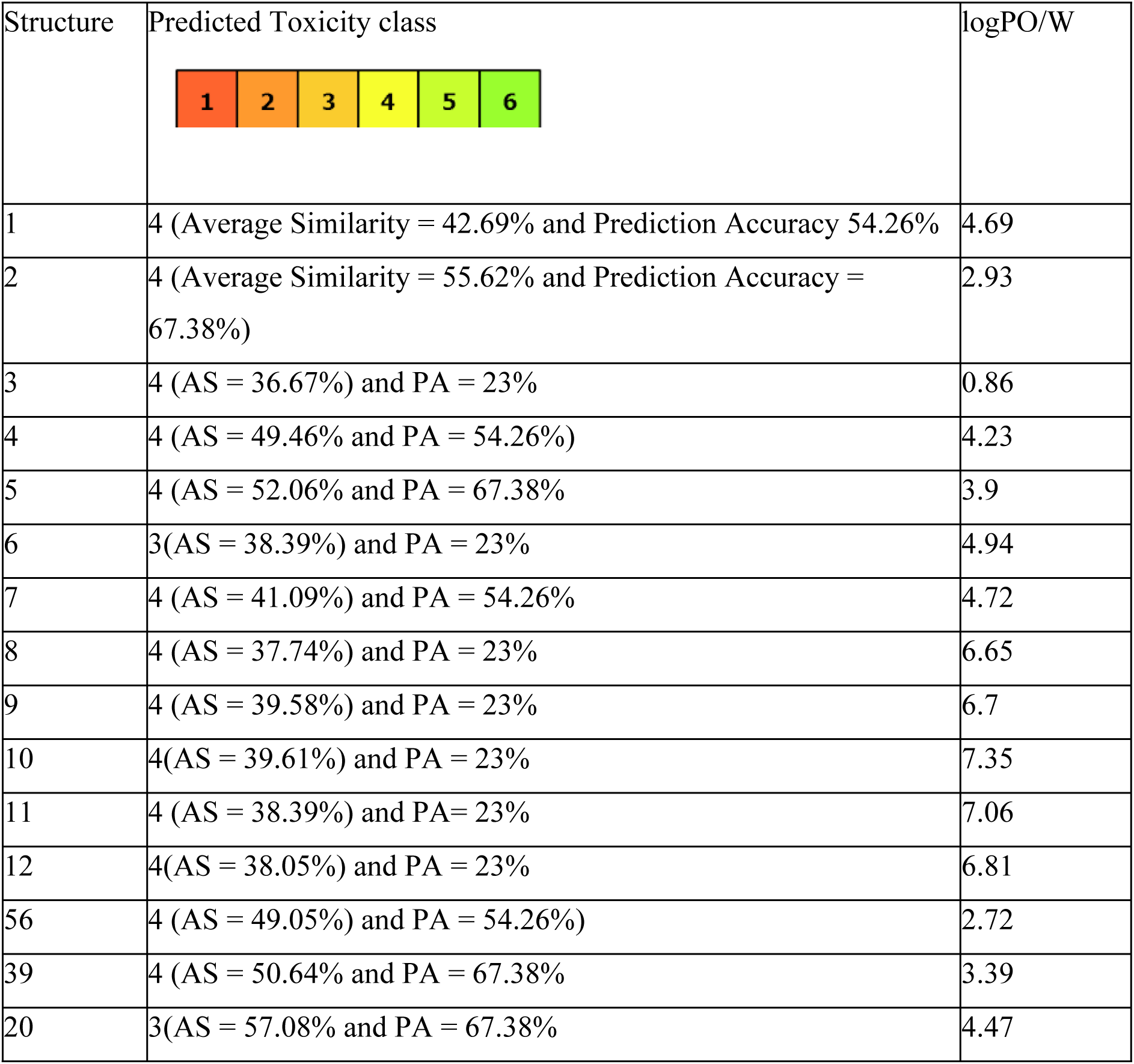
Insilico Toxicity Prediction, PA represent the Prediction accuracy, and AS represents the average similarity.

Table 3: Overall toxicity predictions and log P of the drug molecules

**S20** –CHEMBL ID 1910196 ( 4-[5-(6-hydroxy-1H-indol-2-yl)pyridin-3-yl]benzonitrile)

**S39**- CHEMBL ID 2321945- (2-(7-bromo-2-hydroxy-1H-indol-3-yl)-3-oxoindole-6- carboxylic acid)

**S56**-CHEMBL ID 2321951 (methyl 2-(2-hydroxy-1H-indol-3-yl)-3-nitroso-1H-indole-5- carboxylate)

**S1**- (3,6-diamino-4-(2-chlorophenyl)thieno[2,3-b]pyridine-2,5-dicarbonitrile)

The molecular docking result showed that of all the 12 molecules already reported by Masch et al 2015 (Figure 2), only the co-crystallised ligand **S1** is ranked among the best four molecules that could be investigated further for the treatment of Malaria. It is worth reiterating that all twelve molecules were used as a standard for developing a theory that some of the molecules mined from the CHEMBL database can be repurposed for treating Malaria infection. The molecule labelled **S56. S39** and **S20** were obtained from mining the CHEMBL database using Python coding that we developed.

Table 2 clearly showed that the co-crystallised ligand has a higher binding affinity (s = - 7.1157074 better than **S56** (s=-5.64057302), **S39** (s=-5.42528439) and **S20** (s=-5.26445723), but this did not give it a clear superiority over the remaining three molecules. A closer look at Table 2 above showed that molecule **S56** has a higher number (12 ligand interactions) of ligand interactions within the binding pocket, while **S1** has ten ligand interactions, and molecules **S39** and **S20** have eight ligand interactions. Furthermore, based on the results in Table 2, we cannot say that **S1** behaves better than **S39** and **S20** because, looking at the number of amino acids interacted with in the binding pocket, it is clear that the **S1** molecule interacts with the least number of amino acids in the binding pocket. **S1** interacted with ASP200 and VAL70, while **S56** performed better than all of them, with six different amino acid interactions. **S56** interacted with ILE 562, ASN 186, GLN 185, ARG 141, VAL 70, and THR 138. Of the remaining two ligands, **S39** behaves similarly to **S1**, interacting with only two amino acids VAL 70 and GLN 185, while **S20** perform slightly better than **S1** by interacting with three amino acids, ASN 64, VAL 70 and GLN 185. Therefore, based on the molecular docking results, it might be concluded that **S56** could be successfully repurposed for the inhibition of pfGSK3β, because it competes favorably insilico with the already tested **S1** molecule. Meanwhile, the molecular docking data alone should not be taken as the final bioinformatic data for repurposing a potential drug molecule, therefore, the molecular dynamics simulations of these four molecules were conducted in order to understand the impact of the motion of these molecules in the binding pocket on the stability of the receptor..

### Molecular Dynamics Simulations

Molecular docking results alone could not be used to predict the success of an insilico testing because it does not show the flexibility of the protein during the protein-ligand interaction (Matter et al, 2011 and Kitchen et al, 2004) In this research work, we are focusing on an insilico approach towards the discovery of potential anti-malaria drugs, which could be developed into a new drug moiety for the treatment of malaria. The docking scores have given us a possible drug that performs better than the bioassayed drug molecules (**S1**). To give a deeper insight into the binding behaviour of the biomolecules, Molecular Dynamics Simulations were carried. The molecular dynamics simulations were run on Gromacs 2023.1 using the CHARMM all-atom forcefield, and the Molecular Dynamics Simulations were carried out for 100 ns (50000000 steps). The Root Mean Square Fluctuation (RMSF, Figure 7) and Root Mean Square Deviation (RMSD, Figure 8) trajectory analysis of the MD simulation are results are colour coded as **S20 =** black, S1 = Red, S39 = Green, S56 = blue. Finally, the snapshots of the molecular motion were taken at intervals of 10 ns, 40 ns, 70 ns, 80 ns, 90 ns and at the end of the simulation run(100 ns) (Figures 9,10, 11, 12, 13 and 14 respectively).

**Figure 7:**
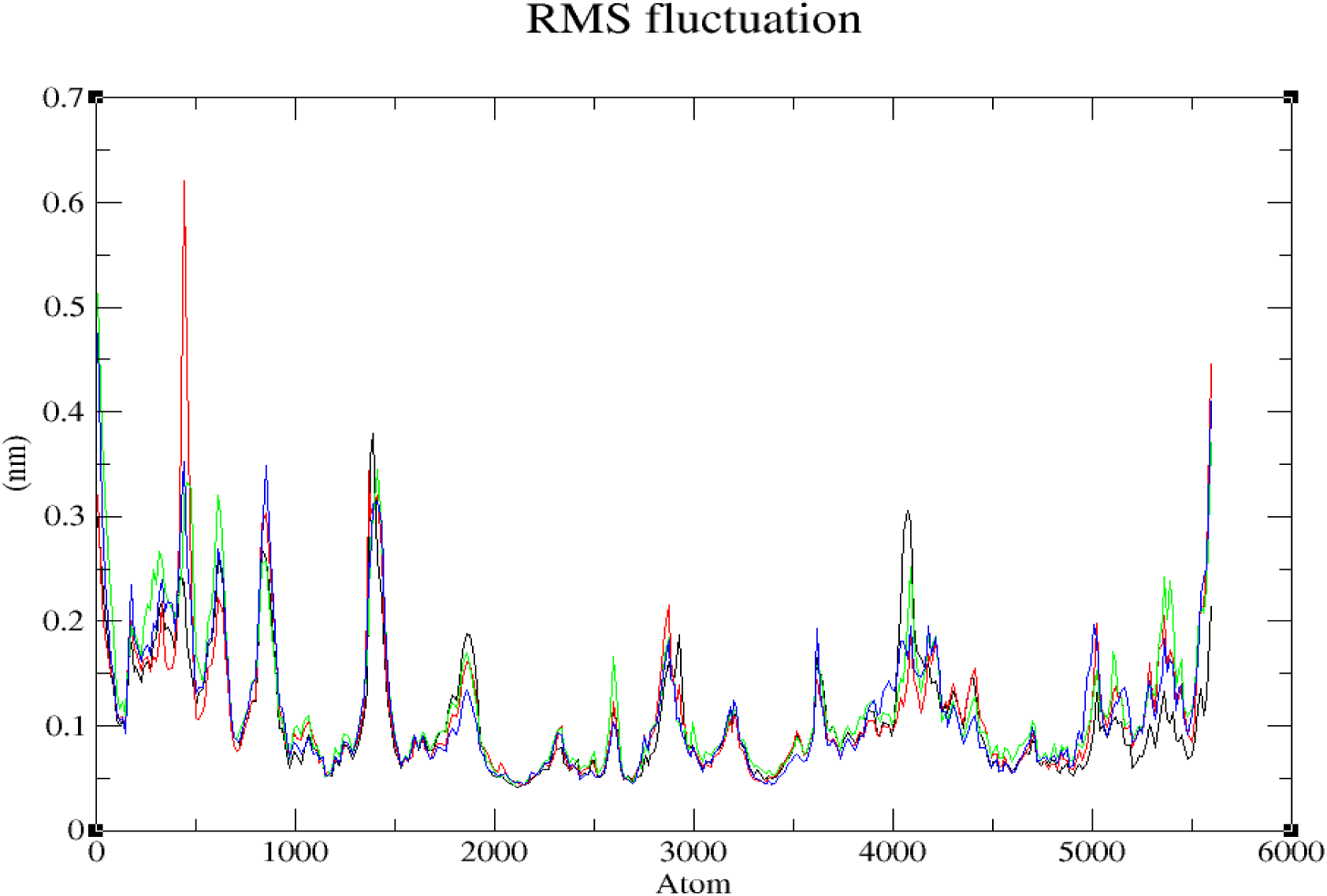
RMSF result of the ligands in the receptor during the simulation.

**Figure 8:**
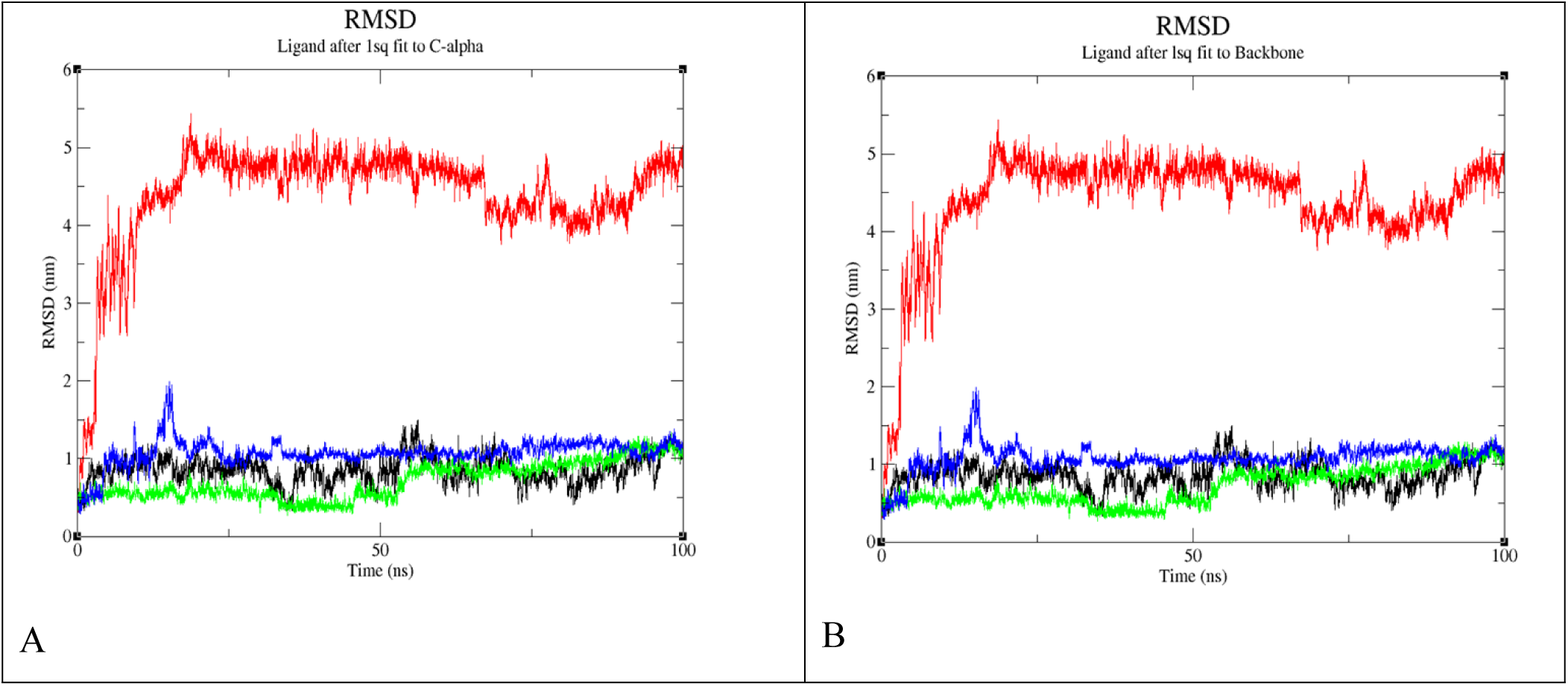
A. RMSD analysis of the ligands in fit to the C-alpha, and B. RMSD analysis of the ligands fit to the backbone.

**Figure 9:**
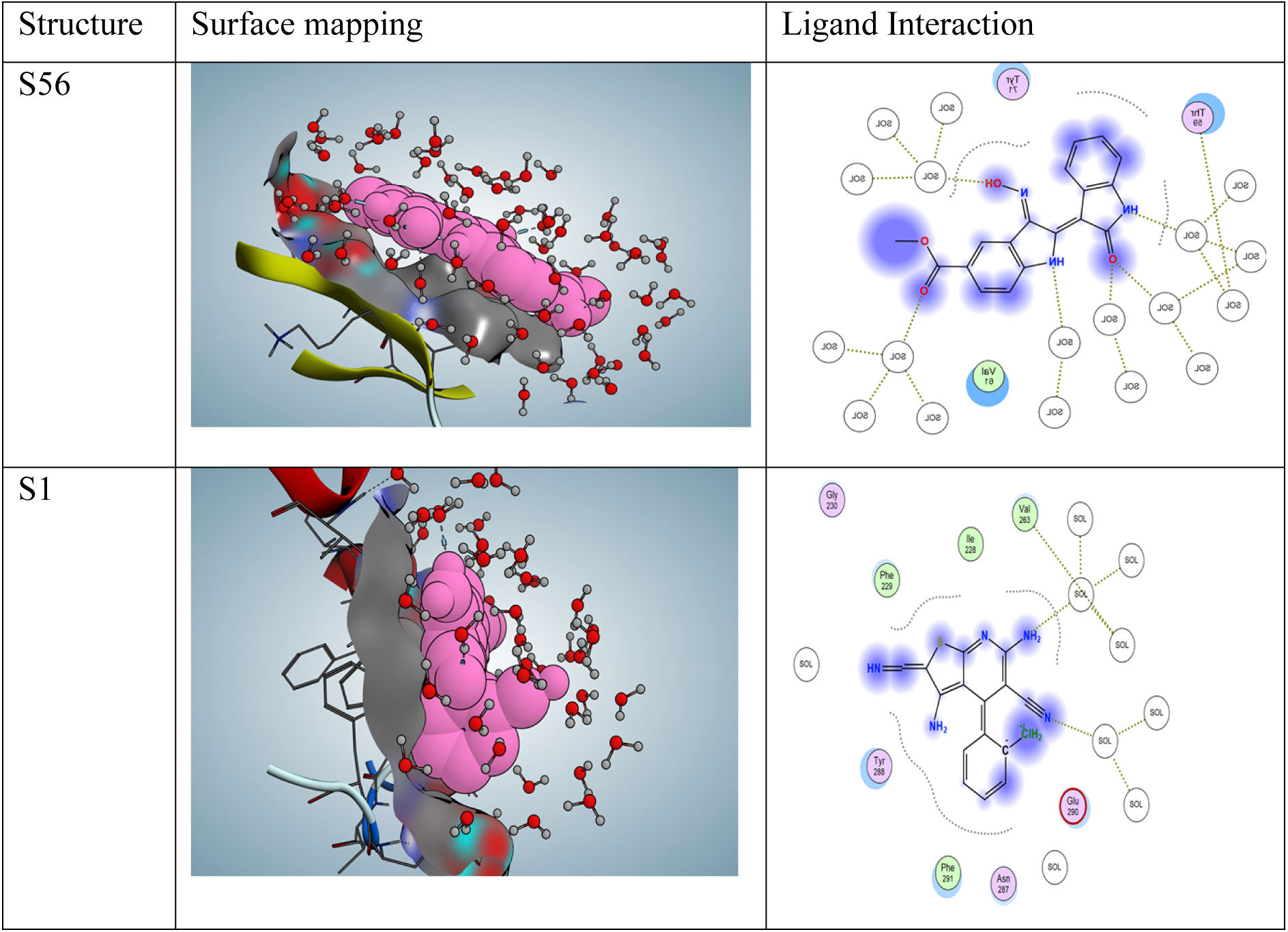
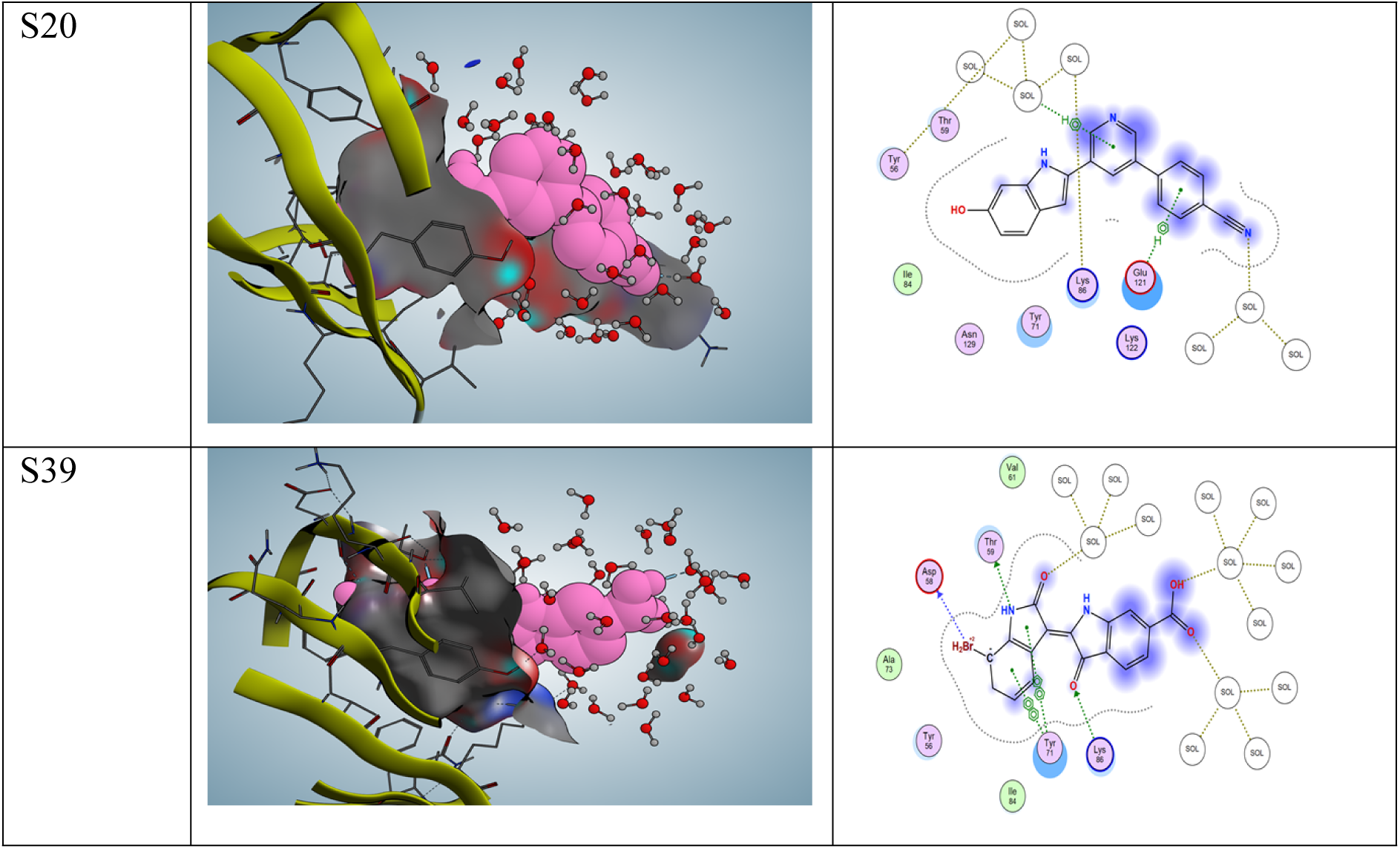
molecular motion at 10 ns

**Figure 10:**
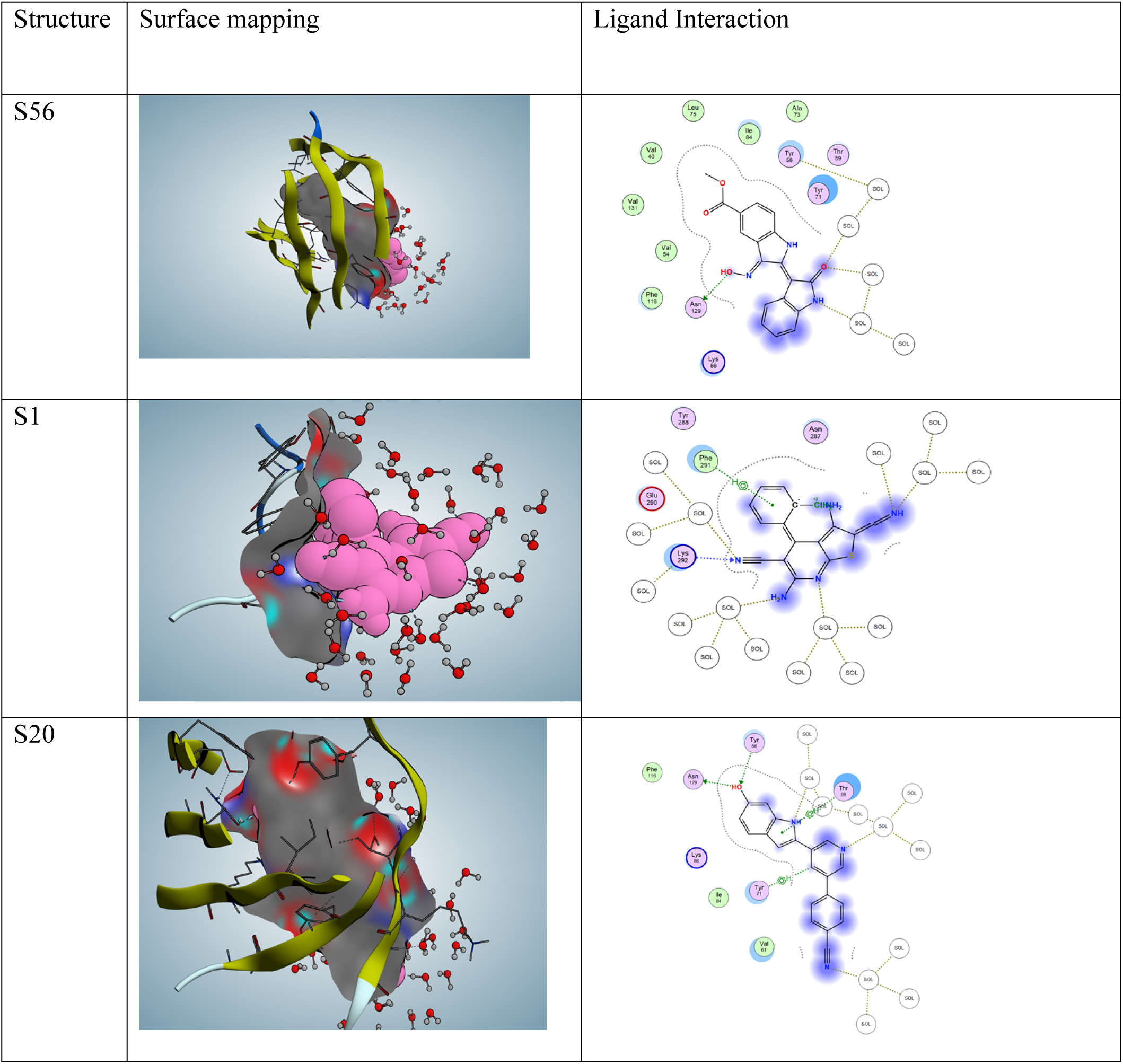
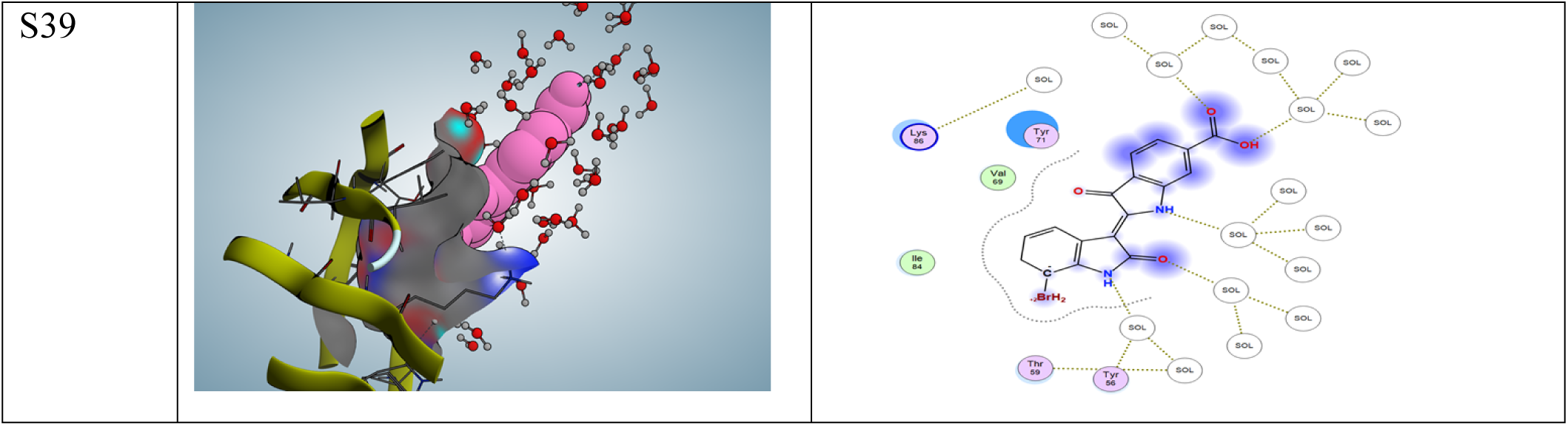
Molecular motion at 40 ns

Root Mean Square fluctuation shows a region in the protein-ligand interaction where the ligand caused a significant change in the protein structure, the instability of the co-crystallised ligand between 0 to 1000 atoms also supported the behaviour that we observed in the RMSD plot(Figure 8). The repurposed drug molecules showed relatively low fluctuation within the binding pocket

The conformational stability of the protein-ligand complex was revealed by the root mean square deviation (RMSD) result of the complex. Figure 8 above shows that the trajectory of the co-crystalised ligand **(S1)** has a very high RMSD when it interacts with the backbone or the c-alpha chain. Whule the repurposed showed a very low RMSD value between 0 and 2 nm and the deviations appeared to be converged at 1.0 nm, the RMSD value is very important because the lower the RMSD the more compact the protein ligand interaction and the better their lock and key process, this better interaction also supported the molecular docking result which gave a better ligand interactions for compound **S56**, **S20** and **S39**, therefore these molecules might show better inhibitory properties than **S1** against pfGSK3β.

Snapshots of simulations were taken at various intervals – 10 ns, 40 ns, 70 ns, 80 ns, 90 ns and 100 ns. These snapshots enable us to understand the behaviour of the ligand within the binding pocket over the simulation steps. According to Marco De Vico et al 2016, snapshots validate the docking results; a bad molecular docking study will be revealed by the Molecular Dynamics Simulations because they are likely to generate a trajectory where the ligand will be seen as leaving the binding pocket, while a good molecular docking result will show the ligand remaining within the binding pocket through out the Molecular Dynamics Simulations production run. This behaviour can also be further visualised by using the RMSF and RMSD diagrams as shown in Figures 7 and 8 above.

### Snapshot at 10ns

**S56** and the co-crystalised ligand **S1** did not interact with any amino acids in the binding pocket, but with the water in the system, **S20**, undergo hydrophobic interactions (pi-H) with GLU 121, and the rest were with solvent in the environment. **S39** Ligand has better interactions with the receptors than all other ligands at 10 ns, it has hydrogen bonding interactions (H-donor) with THR 59 and ASP 58, further hydrogen bonding interaction (H-acceptor) with LYS 86. Furthermore, **S39** Ligand has a hydrophobic interaction (pi-pi) between the 5-ring of the ligand and the 6-ring of the TYR 71, as well as a pi-pi interaction between the 6-membered ring of the ligand and the 6-membered ring of the TYR 71. It is worthy to mention that even though the co crystallized ligands **S1** and **S56** did not interact with any amino acids, in the binding pocket at 10ns time step, this did not make them less effective, Marco Devivo et al 2016 has observed that water molecules have the ability to influence the binding of a ligand to the receptor, water molecules could either support the ligand-interaction or cause the ligand to be rejected from the binding pocket, fortunately all the snapshots at 10 ns time step, the ligands were not rejected in the binding pocket of pfGsk3β.

#### S40 Simulation results

At 40 ns, S56 interacted with ASN 129, while S1 underwent hydrophilic interactions (hydrogen acceptor) with LYS 292. Meanwhile, the six-membered ring of ligand S1 engaged in hydrophobic interactions (π-H) with PHE 291. S20 underwent hydrophobic interactions with the six-membered ring of TYR 71 (H-π), and the five-membered ring of ligand S20 interacted with THR 59 (π-H). Furthermore, S20’s interaction with the receptor was characterised by hydrogen bond donor interactions with ASN 129, while the remaining interactions occurred with the surrounding solvent. S39 does not exhibit any interactions with amino acids at this stage, only with the solvents present in the environment.

### Simulation at 70 ns

At 70ns S56 still undergo hydrophobic interactions using its N2 18 with 6-ring TYR 71 (A) H-pi and also there was another hydrophobic interaction with the ligand 6 membered ring and the six membered ring of TYR 56 (pi-pi), while S1 undergo hydrophilic interaction (H-donor) with ASN 287 and Hydrogen bond acceptor with TYR 288 and 289, as well as GLU 290, S20 made hydrogen bonding interaction (H-donor) with ASN 129, hydrophobic interactions involving the ligand S20 interaction with 6-ring of TYR 71 (H-pi) and pi-H interaction involving the 6-ring of the ligand and LYS 36, while the rest interaction was with the solvent in the environment. S39 undergoes hydrogen bonding interaction (H-donor) with THR 59, hydrophobic interaction (pi-H) with VAL 131 and another hydrophobic interaction (pi-pi)of the 5-membered ring of the ligand and the 6-membered ring of TYR 56

**Figure 11:**
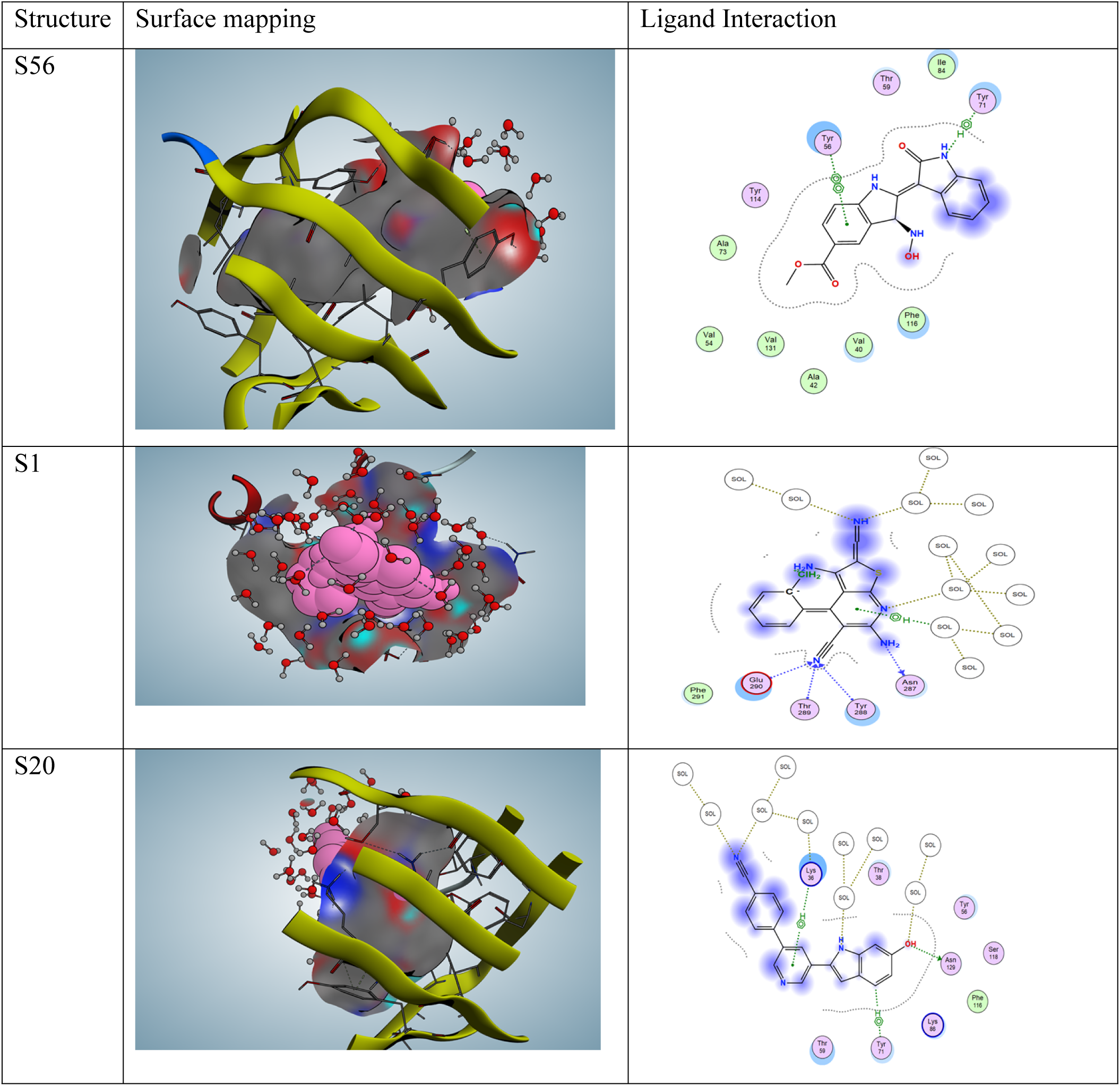
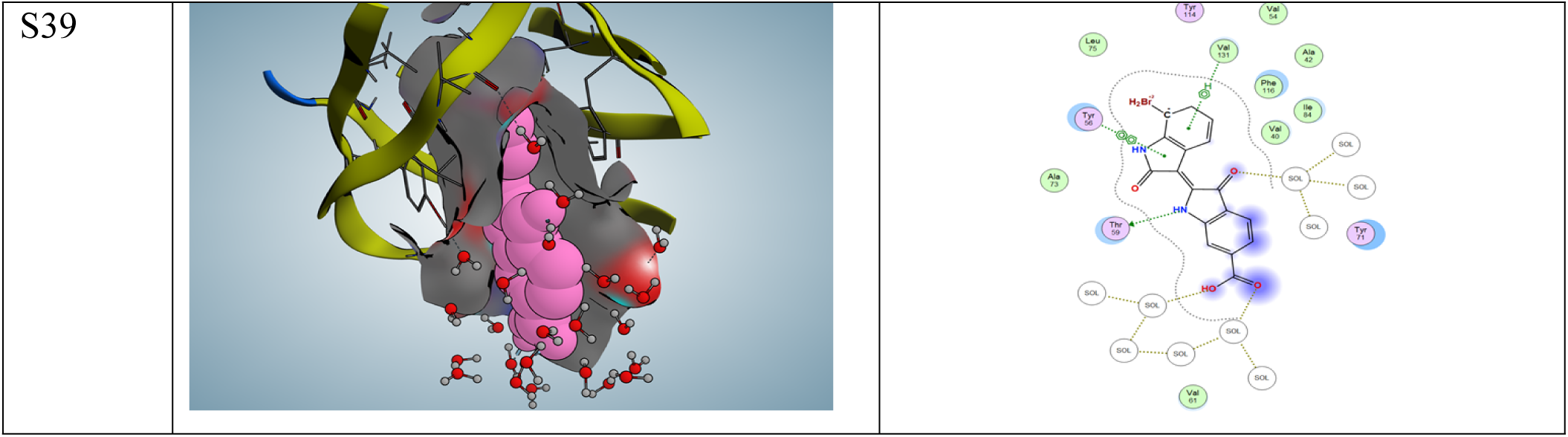
molecular motion at 10 ns

### Simulation at 80 ns

The 6-membered ring of S56 ligand undergoes hydrophobic interaction (pi-pi) with the 6- membered ring TYR 51 of the receptor, S1 only have a Hydrogen bond acceptor with the amino acid ASN 287, while the rest are with the solvent in the environment. S20 ligand only undergoes hydrophobic (pi-H)interactions with the 6-membered ring of TYR 71, while the rest interactions were with the solvents in the environment. S39 ligands only undergo hydrophobic interaction (pi-pi) between the 5-membered ring of the ligand and the 6-membered ring of TYR 56.

**Figure 12:**
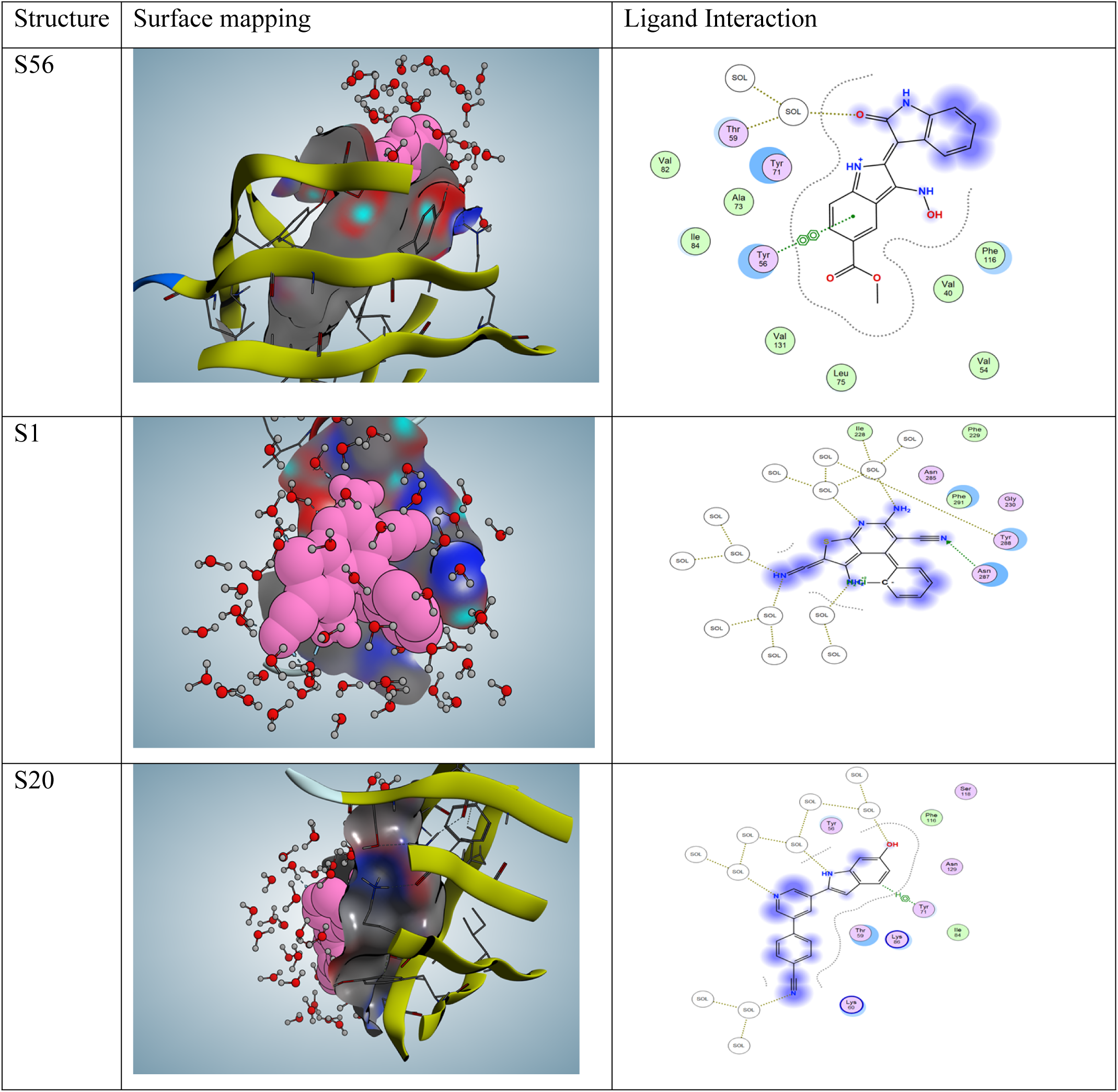
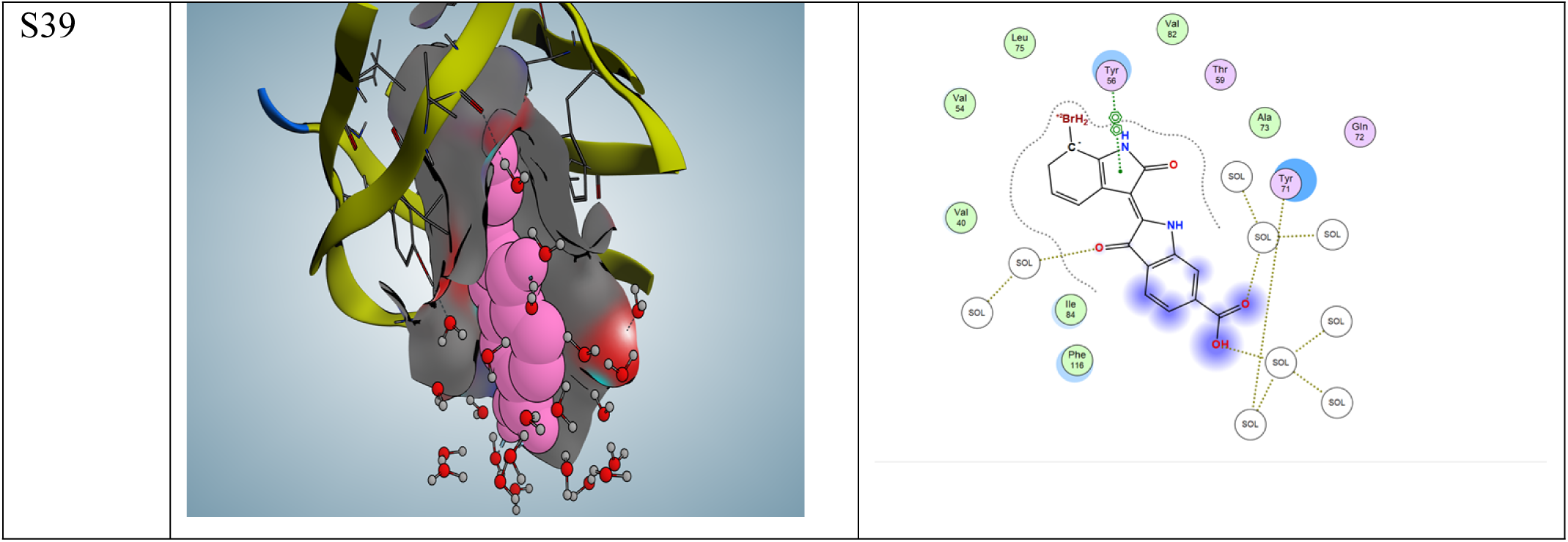
molecular motion at 10 ns

### Simulation at 90 ns

S56 at 90ns did not interact with the receptor but with the water in the environment. At this simulation point, S1 made a hydrogen bond donor with ASN 287, while the remaining were with the water in the environment. S20 undergo hydrogen bonding (H-donor) interaction with ASN 129 and hydrogen bonding (H-acceptor) with TYR56 as well as with solvents in the environment. S39 undergoes a hydrophobic (pi-H) interaction between the 6-membered ring of the ligand and VAL 131, as well as a hydrophobic interaction (pi-pi) involving the 5- membered ring of the ligand and the 6-membered ring of the PHE 116 of the receptor.

**Figure 13:**
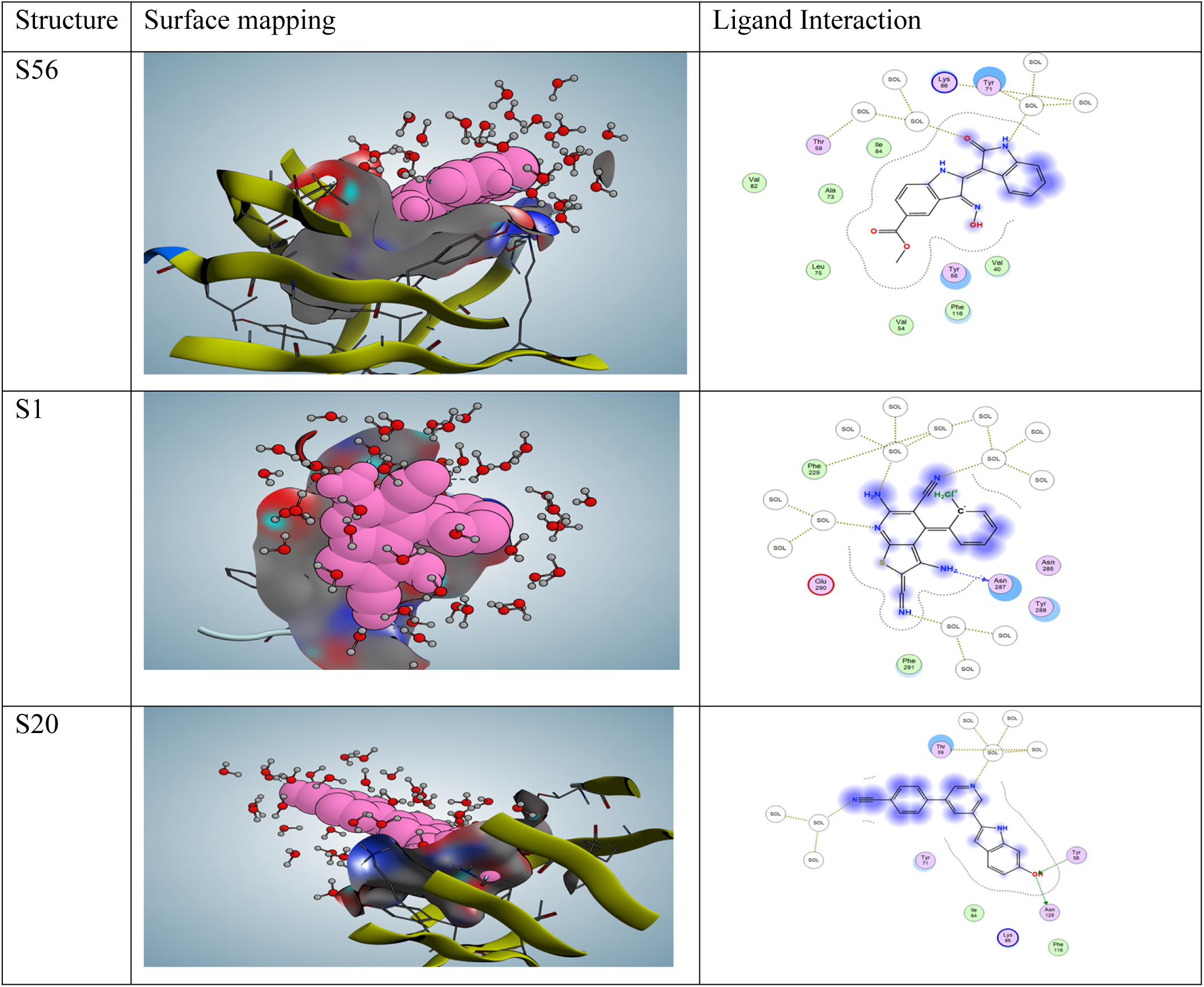
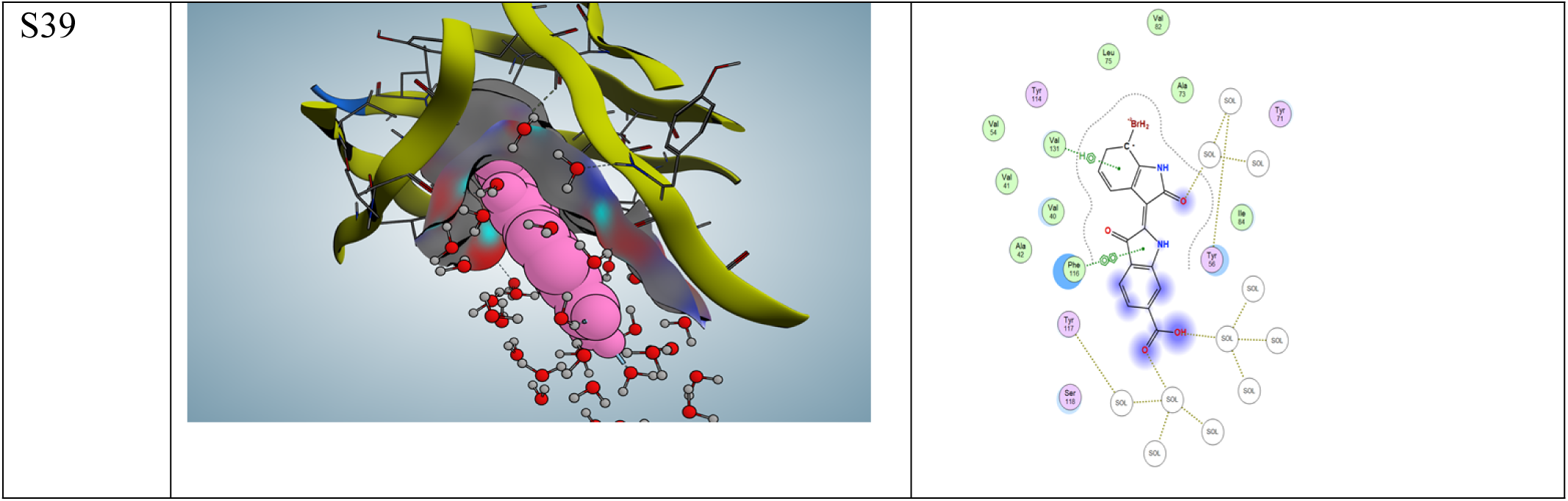
molecular motion at 90 ns

### Simulation at 100 ns

Ligand S56 still maintain a hydrophobic interaction (pi-pi) involving the 6-membered ring of the ligand and TYR56, S1 makes a hydrogen bond interaction with LYS 292, while the rest were with the solvent in the environment. S20 ligands undergo hydrogen bonding interaction (H-donor) with ASN 129 and hydrophobic interaction (pi-H) with THR59; the rest of the interactions were with solvent in the environment. The only amino acid the S39 interacted with at the end of the simulation was a hydrophobic interaction (pi-pi) between the 5-membered ring of the ligand and the 6-membered ring of TYR 56.

**Figure 14:**
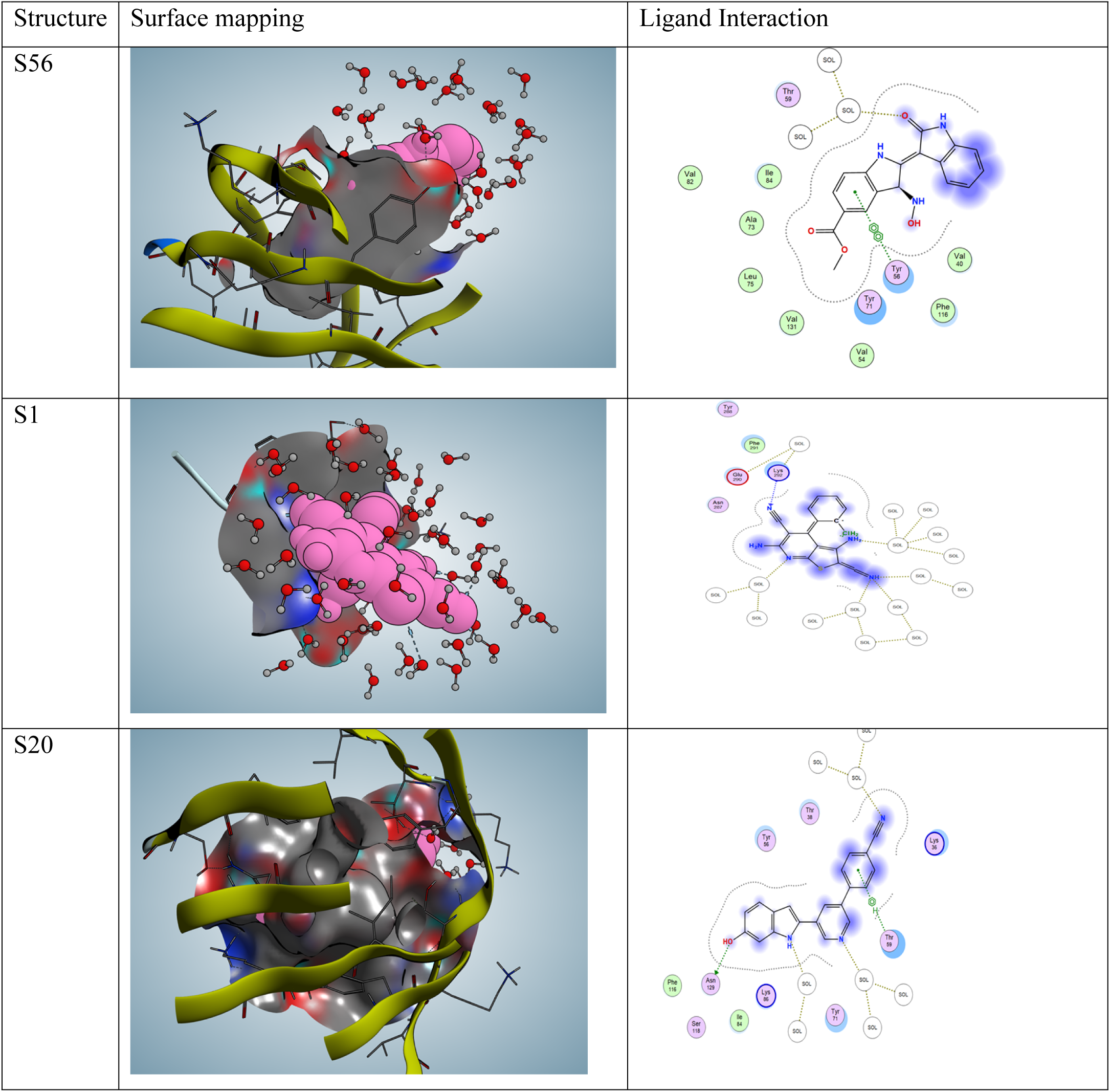
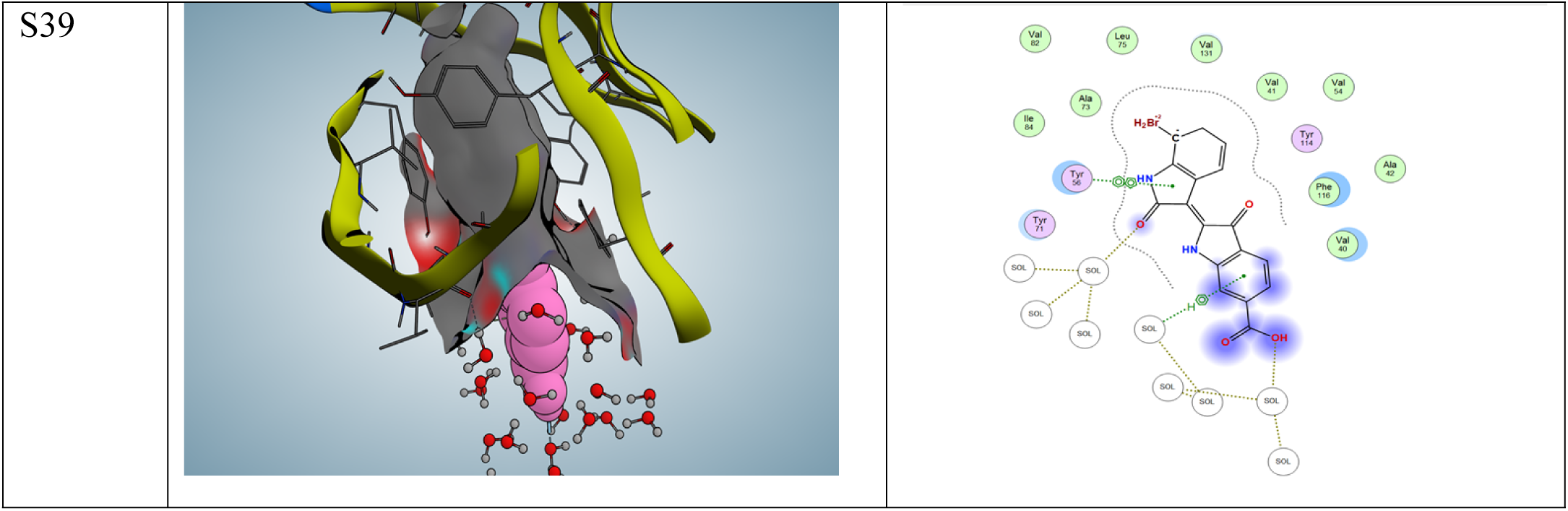
molecular motion at 100 ns

### Insilico Toxicity Profile

In the era of ML (Machine Learning) and the application of Artificial Intelligence towards making research easy and the fact that thousands of laboratory mouse are sacrificed towards testing the toxicity profile of potential drug molecules and only less than 1% of the molecules get to the market, the use of insilico toxicity study could be an important tools towards the study of biological toxicity of new drug molecules and this could reduce the cost that goes into purchasing laboratory rats for toxicological study. The insilico toxicity study was carried out using a web server tool Protox III The reliability and accuracy of this tool have been widely described.(Priyanka et al, 2024)

The predicted overall toxicity showed that all the compounds might be moderately toxic except compound **S6** and **S20** which showed a predicted toxicity of 3, but the toxicity spectrum shown in figure 15 below showed that no single biological molecule is completely cleared, but the spectrum is to indicate what might be expected when carrying out the pharmacokinetics of the potential drug molecules in vivo and invitro.

**Figure 15:**
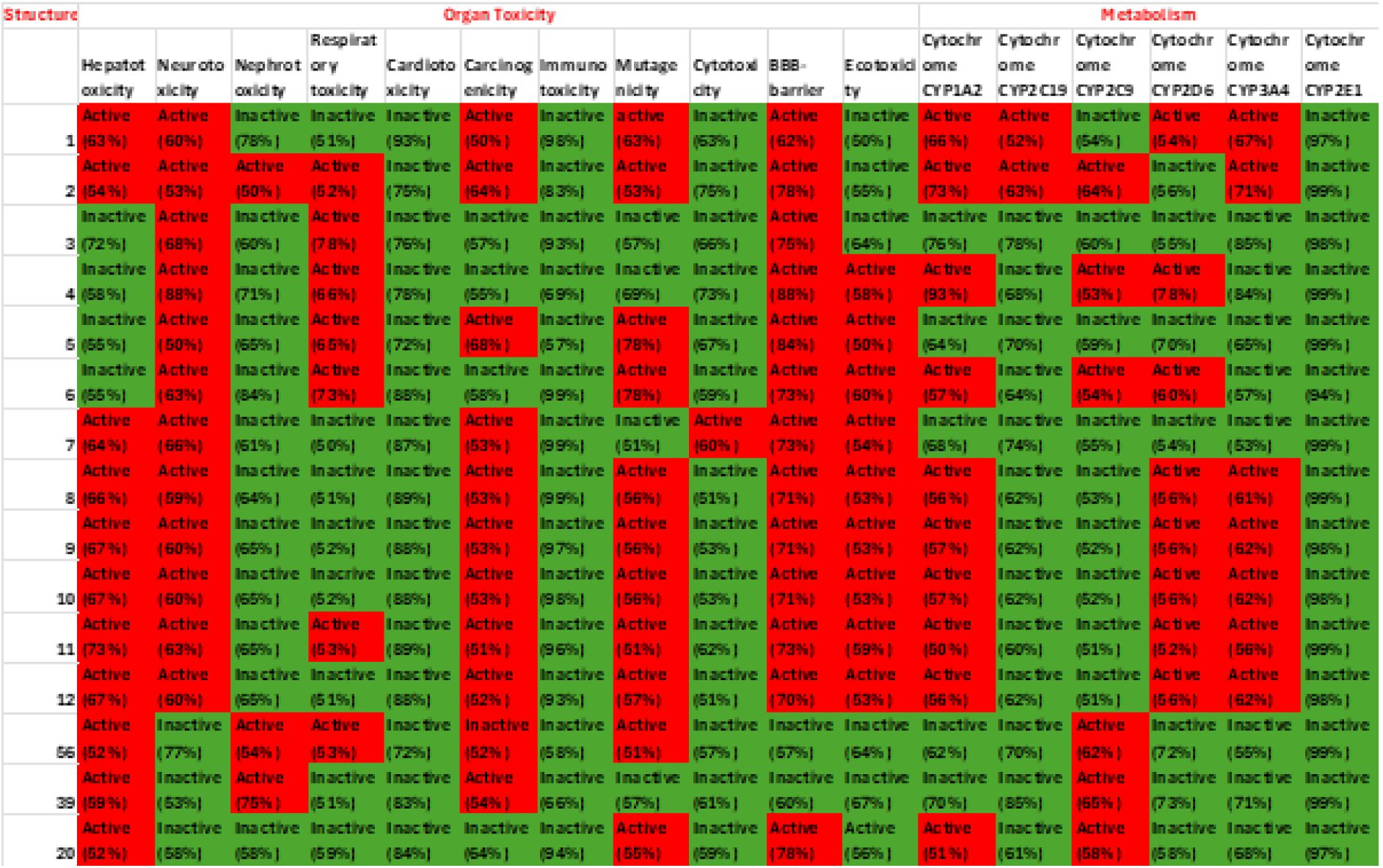
Toxicity profile of the molecules shown in Figure 2 and the best three molecules obtained from the Data mining of the CHEMBL database. The green profile shows that the compounds are inactive in the toxicity category, while the red profile shows that they are active in the toxicity category.

## Conclusion

This research has exploited an insilico approach to repurpose molecules which has been used to inhibit GSK3β found in human beings to inhibit the pfGSK3 enzymes present in Malaria parasites. The ligand interactions of molecules **S56, S39** and **S20** showed good interactions, which are comparable to **S1**; and better than all the other eleven compounds in Figure 2.. Furthermore, the molecular dynamics simulation results showed that these three potential inhibitors of pfGSK3β perform better than the tested molecules (S1) insilico based on the simulation results of RMSF and RMSD, therefore, these drugs could be investigated further for the treatment of Malaria infection..

The insilico toxicity studies of these molecules showed that compounds gave the toxicity spectrum of the molecules, and this will be a useful guide when designing in vivo and in vitro pharmacological studies. According to (Masch & Kunick, 2015), pfGSK-3β (pdb.. 3zdi) is a viable target in the development of novel drugs for the treatment of malaria infection; therefore, we have successfully compiled Python codes which enabled us to mine the CHEMBL database for GSK-3β inhibitors developed for other diseases. This data mining generated 53 molecules, out of which we have been able to obtain three potential molecules (**S56, S39** and **S20**) that could be repurposed for the inhibition of pfGSK-3β enzymes found in the malaria parasite. S56 is the most superior of all these molecules, which is why its structure is shown in the graphical abstract.

## Strengths and limitations

The strength of this study is the use of insilco screening approach, which enabled a rapid and cost-effective evaluation of novel glycogen synthase inhibitors. This computer-based method allowed for the possible identification of drug targets in the treatment of malaria infection. Protein-Ligand molecular docking is a very important tool in understanding the mechanism of Ligand interactions in the receptor by showing us the types of amino acids that are being targeted in order to stop the progression of a disease. This structure-based drug design is successful in the ranking of protein-ligand interactions according to their scores and types of ligand interactions (Maria Batool et al, 2019 and Dongmei Cao et al, 2024), which enables researchers to select the hit molecules for further insilico testing as well as invivo and in vitro experiments. On the other hand Molecular Dynamics simulations have become an important investigational technique to verify some observed experimental data in a non-insilico environment [Van Gunsteren et al, 2008; Laconte et al, 2002; Peter C et al, 2003; Bruschweller et al, 2007; and Markwick et al, 2010] and it is therefore an important insilico technique for repositioning of biological molecules.

Most challenges lie in the fact that an insilico environment might not completely mimic the biological system, as true as this may be, further advances in Machine Learning and Artificial Intelligence make insilico studies an important theoretical approach for predicting the activity of potential molecules, especially in the area of drug repurposing. Furthermore, most challenges that people suggested are firstly the high computational demand of an insilico testing, such as Molecular Dynamics Simulations and secondly the approximation of force fields used in Molecular Dynamics calculations. The issue of High Computational demand has been taken care of by the invention of High Performance Computing system and also the development of desktop and laptop with high performance, this research work was carried out on my high end cheap computer (£4000), having an intel core I9 processor with 24 core and 32 RAM, this laptop have the capacity according to the log file to run 300 ns/day. The second limitation of the appropriate force field has led to researchers incorporating Quantum Mechanical calculations as part of the force field development (Jacob et al, 2011)

## Future work

This study has successfully achieved its aim, and we have utilised an in silico investigation to obtain novel compounds **S56, S20** and **S39**, which, based on the computational study carried out in this research, have the potential of being repositioned as pfGSK3β inhibitors for the treatment of malaria infection. However, further work is necessary to advance and validate these findings, such as *in vitro and invivo* analysis of these molecules before considering taking the repurposed molecules through the phases of clinical trials.

